# A systematic evaluation of the language-of-viral-escape model using multiple machine learning frameworks

**DOI:** 10.1101/2024.09.04.611278

**Authors:** Brent Allman, Luiz Vieira, Daniel J. Diaz, Claus O. Wilke

**Affiliations:** Department of Integrative Biology, University of Texas at Austin, Austin, Texas 78712; Institute for Foundations of Machine Learning, University of Texas at Austin, Austin, Texas 78712

**Keywords:** protein language models, immune escape, machine learning, SARS-CoV-2

## Abstract

Predicting the evolutionary patterns of emerging and endemic viruses is key for mitigating their spread in host populations. In particular, it is critical to rapidly identify mutations with the potential for immune escape or increased disease burden (variants of concern). Knowing which circulating mutations are such variants of concern can inform treatment or mitigation strategies such as alternative vaccines or targeted social distancing. A recent study proposed that variants of concern can be identified using two quantities extracted from protein language models, grammaticality and semantic change. These quantities are defined in analogy to concepts from natural language processing. Grammaticality is intended to be a measure of whether a variant viral protein is viable, and semantic change is intended to be a measure of potential for immune escape. Here, we systematically test this hypothesis, taking advantage of several high-throughput datasets that have become available, and also testing additional machine learning models for calculating the grammaticality metric. We find that grammaticality can be a measure of protein viability, though the more traditional metric ΔΔ*G* appears to be more effective. By contrast, we do not find compelling evidence that semantic change is a useful tool for identifying immune escape mutations.

## 1 Introduction

In an ongoing viral epidemic or pandemic, it is critical to be able to identify variants of concern, that is, viral mutants that may have significant impact on the future course of the epidemic and the potential disease burden caused by it. The gold standard is to perform experimental tests to assess whether mutations confer immune escape *in vitro* [1, 2] or may lead to breakthrough infections *in vivo* [3], but these experiments are labor-intensive, resulting in the characteization of a small number of variants. Larger-scale experiments are possible with deep mutational scanning (DMS) [4–7], yet DMS experiments often require somewhat artificial experimental setups that don’t necessarily reflect the infection conditions viruses encounter in their natural hosts.

As an alternative to experimental strategies, several groups have employed modeling approaches, either by predicting the fitness of variants from epidemiological data [8] or by employing biophysical or other mechanistic modeling approaches to predict the fitness of a mutation from its context in the viral protein and the expected interactions with antibodies and host receptors [9]. The downside of the epidemiological approach is that it is strictly backwards looking and cannot make any prediction for newly emerging variants. By contrast, mechanistic and machine learning (ML) modeling approaches are forward-looking but tend to require intensive compute and extensive experimental data for model calibration.

An ideal modeling approach would be able to make predictions for novel mutations while requiring little or only easily obtainable experimental data. In a recent paper, Hie *et al*. [10] suggested that this goal could be achieved with deep learning models, and specifically with protein language models (pLMs) trained on viral sequence data. Specifically, they argued that they could extract two relevant quantities from the pLMs, the grammaticality of a mutation and the amount of semantic change induced in the viral protein. The concepts of grammaticality and semantic change are borrowed from natural language processing. In the context of viral evolution, they represent whether the mutation can be expressed in principle (grammaticality) and to what extent it may change the interaction of the expressed viral protein with its environment (semantic change). In particular, the core idea of Hie *et al*. [10] was that mutations with large predicted semantic change should be more likely to disrupt the protein–protein interface with neutralizing antibodies, and if those mutations also have high grammaticality, they can be selected for immune escape by evolution *in vivo*. Hie *et al*. [10] considered three different viral systems, SARS-CoV-2, influenza A, and HIV-1, and demonstrated some association between escape mutations and mutations with both high grammaticality and high semantic change. However, since the original publication of Hie *et al*. [10], their approach has not seen much application or systematic evaluation (but see [11]). Thus, whether the language modeling approach is useful to predict variants of concern remains unclear.

Here, we take advantage of several high-throughput experimental datasets that have been published since Hie *et al*. [10] and ask how well the concepts of grammaticality and semantic change correlate with experimentally validated immune escape mutations. We further ask to what extent the concept of grammaticality relates to more conventional concepts such as protein stability, and whether there are alternative modeling approaches that can predict the viability of mutations more reliably. Overall, while we find that both grammaticality and semantic change are somewhat informative about escape mutations, neither is sufficiently predictive to provide a compelling use case. In particular, for the purpose of distinguishing viable from inviable mutations, we find ΔΔ*G* values to be more effective than grammaticality. For the purpose of identifying mutations with immune escape potential, we don’t find that semantic change is sufficiently discriminatory, but we also are not aware of any other method or metric that would perform better. Overall, we conclude that the task of predicting likely variants of concerns with current pLM frameworks is inadequate and requires extensive future work and potentially different approaches.

## 2 Results

Before presenting new results, we first provide an overview of the fundamental concepts, datasets, and models used in our analysis. The model architectures used in the analysis are key for contextualizing how we address whether grammaticality and semantic change are useful for interpreting the evolution of antigenic escape.

### 2.1 Background and model setup

Protein language models take principles from natural language processing and apply them to protein sequences. Instead of operating on sequences of words, however, protein language models operate on sequences of amino acids. But apart from this difference, model architec-tures and modeling principles carry over nearly one-to-one. Where natural language models are trained to predict masked tokens in a sentence (i.e. BERT-style self-supervised learning) or the next token in a sentence fragment (i.e. GPT-style self-supervised learning), protein language models are similarly trained to predict masked amino acid tokens (e.g., ProtBERT [12] and ESM [13]) or the next amino acid token (e.g., ProGen [14]). Therefore, it makes sense to assess to what extent other concepts from natural language processing also translate to protein language models.

In natural language processing, it has been fruitful to distinguish the concepts of grammaticality and semantics of words [15, 16]. Grammaticality indicates that a word fits in its location in a sentence purely based on the rules of grammar (for example, the word is a noun and a noun is required at its location in the sentence) whereas semantics describes the meaning of words. A word can have high grammaticality in a sentence but be a poor fit semantically or vice versa. In well-formed sentences each word has high grammaticality and good semantic fit. Hie *et al*. [10] built on these concepts and proposed that they could be applied to protein language models, and more specifically to evaluate variants of viral proteins. The grammaticality of a variant should be related to whether the mutation can be made in principle, i.e., whether the mutated protein still can be expressed and fold, and the semantic change of a variant should represent to what extent biological function has changed (Fig. 1). For example, a mutation on the surface of a viral spike protein with high semantic change is more likely to result in structural changes that disrupt protein–protein interfaces and potentially enable immune escape. Since the concepts of grammaticality and semantic change are derived from natural language modeling, in the following we will refer to them in aggregate as the “language-of-viral-escape model.”

**Figure 1:**
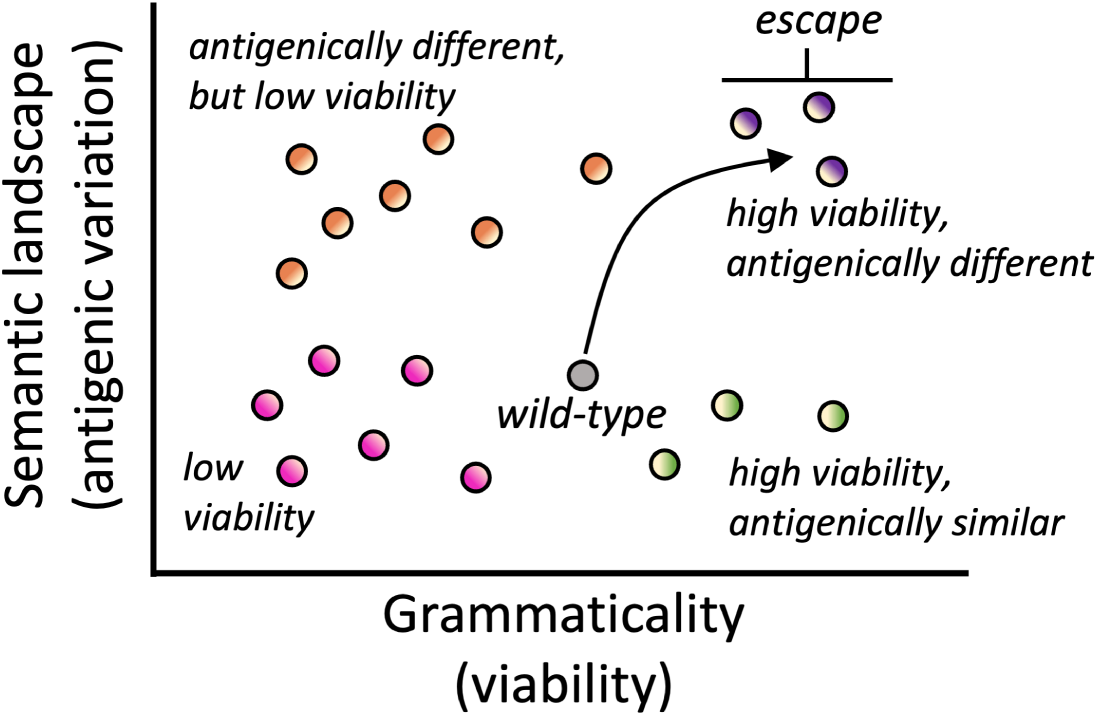
Mutations placed on a wild-type genetic background can have impacts on both viability and antigenic variation. In the language-of-viral-escape model [10], grammaticality is an analogy for protein viability and semantic change is an analogy for change in the surface properties of the spike protein and thus specifically antigenic variation. The hypothesis is that the most effective antigenic escape mutations will be both highly grammatical and semantically different. Schematic drawing modified from Hie *et al*. [10].

However, we caution that the analogies employed by the language-of-viral-escape model are imperfect, and the two quantities grammaticality and semantic change may not ade-quately quantify a viral protein’s viability and immune escape propensity. For example, antigenic escape is just one possible phenotype that can result from mutational changes on a surface antigen. Viral spike proteins need to bind host receptors for cell entry, and *a priori* it is not clear why a large semantic change should specifically correspond to weakened interactions with an antibody but not to weakened interactions with the host receptor. Immune escape variants need to disrupt the binding of neutralizing antibodies while preserving binding to the host receptor. Similarly, even if grammaticality as defined by Ref. [10] is associated with protein viability, there may be alternative ways of calculating grammaticality with more biophysically-based quantities, such as ΔΔ*G*, which better capture viability.

### 2.2 Alternative datasets and alternative models to validate the language-of-viral-escape model

Hie *et al*. [10] trained three bespoke models for three different viral systems, influenza hemagglutinin (influenza HA), HIV-1 envelope glycoprotein (HIV Env), and SARS-CoV-2 spike protein. At the time of their publication, mutational data for SARS-CoV-2 was sparse. To expand on this work, we benchmarked their model on several more comprehensive SARS-CoV-2 datasets that have since become available [5–7]. Additionally, we compared these results to data from influenza HA [4, 17] and HIV Env [18, 19] to assess the generalization of language-of-viral-escape to other viral systems.

For SARS-CoV-2, Hie *et al*. [10] validated their semantic change and grammaticality scores on only 12 escape mutations and 16 non-escape mutations reported by Baum *et al*. [20]. In a direct comparison of these 28 mutations, the escape mutations had consistently higher semantic change than the non-escape mutations and both groups of mutations had high grammaticality (Fig. 2A, inset). However, when calculating grammaticality and semantic change for a much larger DMS dataset [7], we found that escape mutations were not enriched for high semantic change and many escape mutations had surprisingly low grammaticality scores (Fig. 2A).

**Figure 2:**
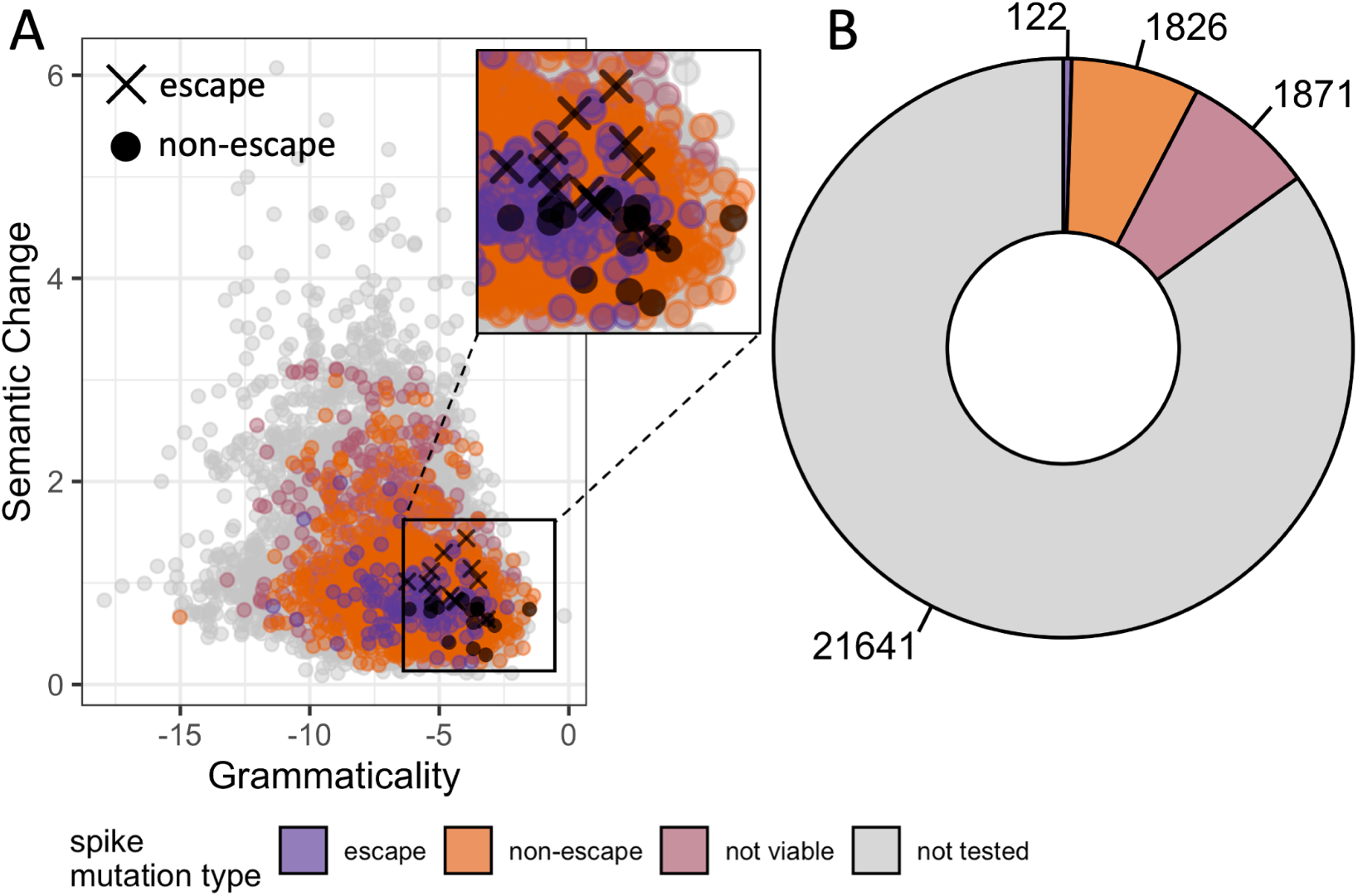
(A) Semantic change and grammaticality scores of mutations predicted by Hie *et al*. [10] attempt to provide insight into antigenic escape. (B) Deep mutational scanning data [7] validate propensity for a mutation to escape antibody pressure, but also give insight into the viability of mutations. Purple represents mutations that had an escape fraction greater than 0.5 in DMS experiments. Orange represents mutations that had an escape fraction below 0.5. Pink represents mutations that render the virus not viable. Gray represents mutations that were not tested for antigenic escape. Black X’s and circles are escape and non-escape mutations verified by Baum *et al*. [20], respectively.

These observations suggested that a more systematic analysis of grammaticality and semantic change in relation to the measured phenotypes of viral proteins was needed. Since the DMS data provided us with measurements for both viability and immune escape for all tested mutations, we could systematically investigate the association of grammaticality and semantic change with respect to both phenotypes. And even though DMS data covered only a fraction of the total mutation landscape for SARS-CoV-2, it increased the number of mutations available for validation by over two orders of magnitude (Fig. 2B).

In addition to this more systematic validation, we also asked whether there are alter-native approaches to calculating grammaticality. For example, the machine learning-guided protein engineering community develops deep learning frameworks with the express purpose of predicting allowed mutations, in particular structure-based frameworks [21–24]. Addi-tionally, virtually all protein language models have been trained on masked token prediction and should also be able to predict whether a mutation is permitted or not [13, 23, 25–27]. Finally, changes in protein free energy, measured by ΔΔ*G*, have long been used to assess whether a mutation at a given site in a protein is permitted or not [28]. All of these model-ing approaches could potentially yield measures of grammaticality that are more informative than the one proposed by Hie *et al*. [10].

To evaluate the effectiveness of grammaticality, we employed a variety of deep-learning models (Table 1). First, we leveraged two protein language models: the bidirectional long short-term memory (BiLSTM) model trained by Hie *et al.* [10] and the transformer-based ESM2 model from Meta [13]. The bespoke BiLSTM was trained on specific SARS-CoV-2, influenza, and HIV protein sequence datasets for their respective surface antigens. ESM2 was trained on the UniRef50 dataset (*∼*65M sequences clustered at a 50% sequence similarity) [29]. Furthermore, for ESM2, we employed two distinct approaches of calculating grammaticality. The first, which we call “unmasked,” was proposed by Hie *et al.* [10]. For each possible mutation at a site, this approach makes a separate prediction, which is then interpreted as grammaticality score. The second, which we call “masked”, is canonical BERT-style masked language modeling (MLM): mask the focal site and use the sequence context to predict amino acid propensities for all twenty amino acids at that site. We convert these propensities into grammaticality scores.

**Table 1:** Models used to calculate the grammaticality of mutations.

|  | Input Data |  | Training Mode |  | Model Outputs |  |
| --- | --- | --- | --- | --- | --- | --- |
| | Sequences | Structures | Self-supervised | Supervised | AA propensities | $\Delta\Delta G$ |
| BiLSTM (Hie et al.) | ✓ |  | ✓ |  | ✓ |  |
| ESM2 | ✓ |  | ✓ |  | ✓ |  |
| MutComputeX |  | ✓ | ✓ |  | ✓ |  |
| MutRank | ✓ | ✓ | ✓ |  | ✓ |  |
| Stability Oracle |  | ✓ |  | ✓ |  | ✓ |

Second, we utilized the structure-based machine learning models MutComputeX [21], MutRank [22], and Stability Oracle [30]. MutComputeX uses a self-supervised 3-dimensional residual neural network (3DResNet) architecture trained on millions of spatially represented protein microenvironments to predict the masked amino acid using the cross entropy loss. MutRank is a two-stage self-supervised graph-transformer framework trained to learn protein representations tailored for protein engineering applications. Using a masked microenvironment as input, MutRank is first trained to predict the masked amino acid from its microenvironment, akin to MutComputeX, and then, with “From” and “To” amino acid CLS tokens as additional inputs, it is self-supervised fine-tuned with a rank loss to predict mutational propensities (i.e. predict the EvoScore) based on the site-specific amino acid distribution obtained from a protein’s multiple sequence alignment. For a particular residue, the EvoScore can be predicted for mutating from the wildtype amino acid to all 20 amino acids, providing amino acid mutational propensities. Amino acid propensities for both self-supervised models are converted into grammaticality scores as described [10]. Finally, Stability Oracle uses the same architecture and training procedure as MutRank. However, rather than being self-supervised fine-tuned to predict the MSA-based EvoScore, it is supervised fine-tuned on a curated subset of the mega-scale ΔΔ*G* cDNA dataset [31] that is augmented with thermodynamic permutations (TP), cDNA117K + TP. Thus, rather than predicting amino acid mutational propensities, Stability Oracle predicts ΔΔ*G* values (kcal/mol) for mutations, where more negative ΔΔ*G* values are considered more grammatical.

For all models, we rank-ordered grammaticality scores and then normalized the ranks by the total number of mutations to arrive at a relative rank score. This score, which is a value between 0 and 1, allowed us to directly compare predictions from widely differing models producing outputs on different scales and with differing distributions.

### 2.3 Different grammaticality measures can distinguish between viable and nonviable mutations

We calculated six different grammaticality scores for all possible point mutations in the SARS-CoV-2 spike protein, using the five different models described above (Table 1) and employing two separate strategies for ESM2, masked and unmasked. We had available viability and immune escape data from Starr *et al*. [7] for *∼*15% of these mutations (Fig. 2B). We found that with the exception of the unmasked ESM2 model, all models produced significantly higher grammaticality scores for viable mutations than for nonviable mutations (Fig. 3A). Among the language models, the bespoke model by Hie *et al*. [10] displayed a larger effect size than the masked ESM2 model. However, in both cases, while the difference in grammaticality scores among viable and nonviable mutations was significant, the effect size was small and the interquartile ranges of the two distributions showed extensive overlap. Thus, a high or low pLM grammaticality score was not predictive of whether a mutation would be viable or not, respectively.

**Figure 3:**
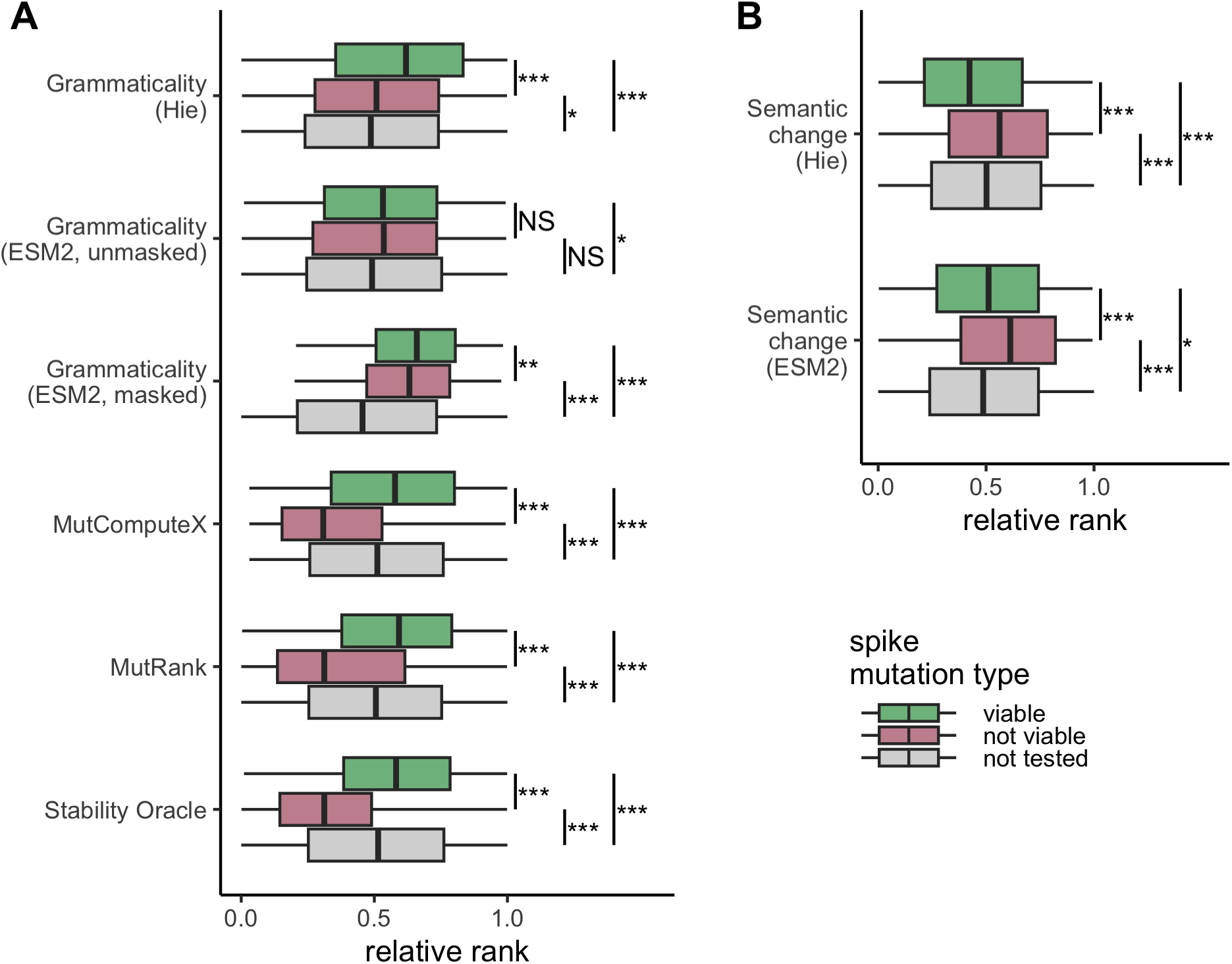
All possible mutations of the coronavirus spike protein DMS experiment [7] tested under different models. Colors represent mutations that are viable (green), not viable (pink), or not tested (grey) in the DMS experiment. The values predicted for each mutation are ranked and then normalized to be between 0 and 1. (A) Grammaticality scores for each of the six models. Note that the ranks for Stability Oracle are reversed since negative ΔΔ*G* values represent more stable mutations. (B) Semantic change scores for both the Hie *et al*. [10] model and the ESM2 model. Results of the Mann-Whitney U rank test are indicated as: NS *→* not significant, *∗ → p < .*05, *∗ ∗ → p < .*01, *∗ ∗ ∗ → p < .*001.

We saw more substantive differences among the three structure-based frameworks: MutComputeX, MutRank, and Stability Oracle. For all three models, the median grammaticality score for nonviable mutations fell into the bottom quartile of grammaticality scores for viable mutations (Fig. 3A). While effect sizes among the three models were somewhat comparable, overall the strongest separation was seen for Stability Oracle, which predicts ΔΔ*G* values rather than amino acid propensities. These results suggest that changes in protein stability, a biophysical property, best capture changes in protein viability, as compared to grammaticality scores computed from amino acid propensities from sequence-or structure-based self-supervised models.

We also assessed how semantic change related to protein viability. We calculated semantic change for two models, the bespoke model by Hie *et al*. [10] and the generic model ESM2. Structure models work at the microenvironment level and do not generate representations for the entire protein, which makes them not suitable for semantic evaluation. Results were comparable for both models (Fig. 3B). Semantic change on average was smaller for viable mutations than for nonviable mutations. This result makes intuitive sense—a mutation that changes more of the protein biochemistry (creates a larger semantic change) should also be more likely disruptive to the protein and render it nonviable. However, this result puts into question whether grammaticality and semantic change are truly independent quantities. In fact, in Fig. 2A we can see a weak negative correlation between grammaticality and semantic change (Pearson’s r = *−*0.26, p-value *<* 10*^−^*^15^).

Finally, we repeated these analyses for two additional viral surface proteins: the hemagglutinin (HA) of influenza A virus (Figs. S1 and S3) and the envelope glycoprotein (env) of human immunodeficiency virus (HIV) (Figs. S2 and S4). Results were generally consistent with what we had seen for SARS-CoV-2 spike. All grammaticality scores except those calculated with ESM2 unmasked were on average larger for viable mutations than for nonviable mutations (Fig. S3A, Fig. S4A). However, the structure-based methods and in particular Stability Oracle tended to perform better than the bespoke model by Hie *et al*. [10]. And again, viable mutations tended to have lower semantic change than nonviable mutations, regardless of the model according to which semantic change was calculated (Fig. S3B, Fig. S4B).

### 2.4 Semantic change does not predict immune escape

We next proceeded to assess the relationship between semantic change and the immune escape phenotype, which was also assessed in the various DMS experiments [4, 7, 17–19]. Notably, for SARS-CoV-2 spike, we found no significant difference in semantic change among escape and non-escape mutations (Fig. 4A). This result was independent of the model we used to calculate semantic change. We saw the same result for HIV env (Fig. S5A). Only for influenza virus HA was there a significant (but small) difference in semantic change between escape and non-escape mutations, and only for the bespoke model trained by Hie *et al*. [10] (Fig. S6A). In aggregate, these results suggest that semantic change is not a reliable or strong predictor of immune escape, even when trained on mutant data for a particular viral protein. For completeness, we also investigated whether grammaticality scores differed among escape and non-escape mutations. Most models showed no significant difference or only minor differences for SARS-CoV-2 spike (Fig. 4B). A few more models displayed significant differences for HIV env (Fig. S5B) and influenza virus HA (Fig. S6B). In the latter case, the bespoke model by Hie *et al*. [10] showed immune escape mutations to be substantially more grammatical than non-escape mutations. Notably, this is the opposite prediction from SARS-CoV-2 spike, where the bespoke model predicted escape mutations to be less grammatical. In aggregate, these results reiterate that grammaticality scores are not necessarily orthogonal to whether or not mutations are escape mutations, and that in general grammaticality and semantic change are somewhat confounded with each other.

**Figure 4:**
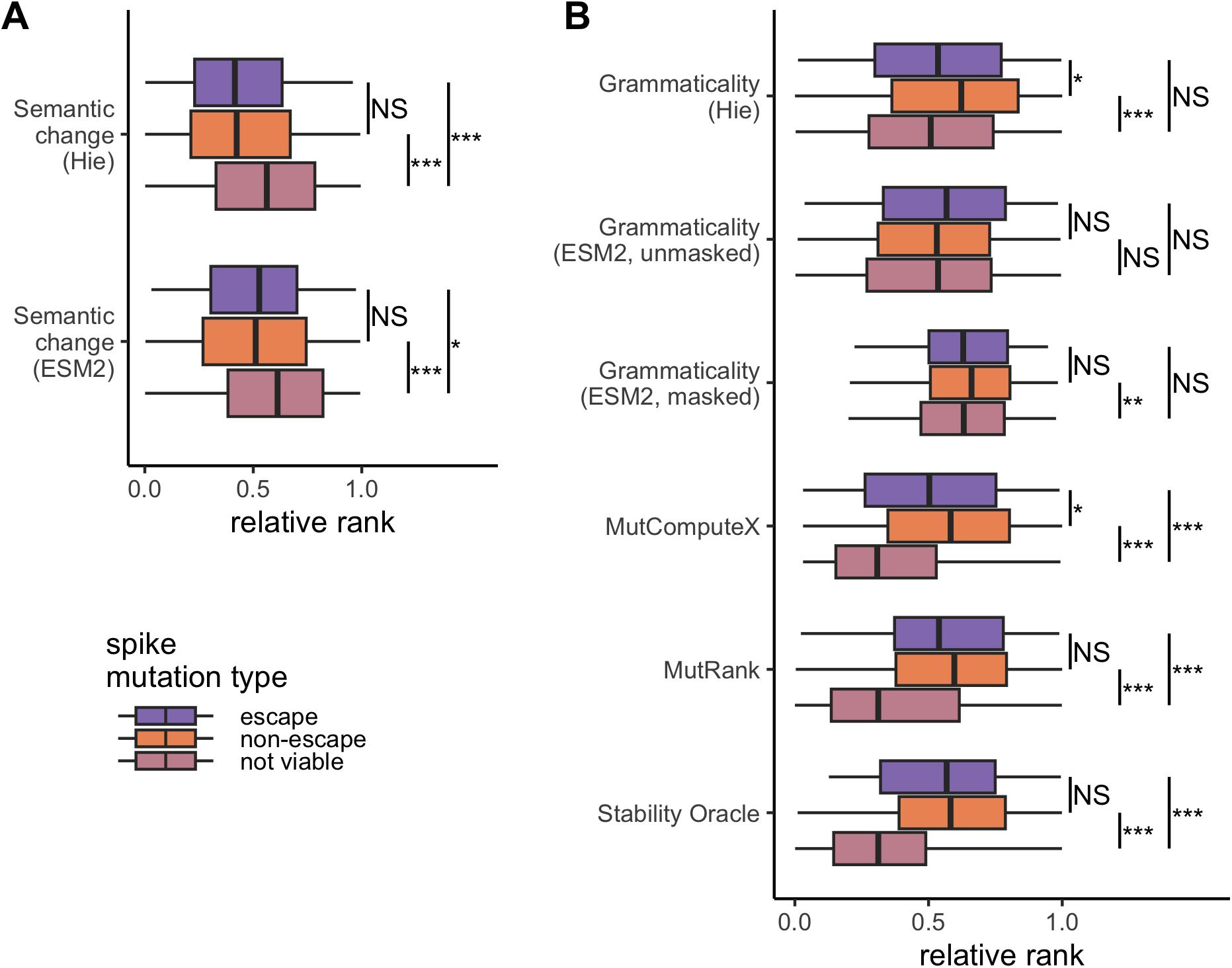
All possible mutations of the coronavirus spike protein DMS experiment [7] tested under different models. Colors represent mutations that confer escape (purple), non-escape (orange), or not viable (pink) in the DMS experiment. The values predicted for each mutation are ranked and then normalized to be between 0 and 1. (A) Semantic change scores for both the Hie *et al*. [10] model and the ESM2 model [13]. (B) Grammaticality scores for each of the six models. Note that the ranks for Stability Oracle are reversed since small ΔΔ*G* values are consistent with higher stability. Results of the Mann-Whitney U rank test are indicated as: NS *→* not significant, *∗ → p < .*05, *∗ ∗ → p < .*01, *∗ ∗ ∗ → p < .*001.

### 2.5 Semantic change is weakly correlated with antibody binding

Instead of considering mutations that have been classified into two categories, escape or non-escape, it may be more useful to ask whether semantic change correlates with the strength of antibody binding for different protein variants. We asked this question on a more comprehensive dataset of 32,768 variants of the SARS-CoV-2 spike protein for which binding constants have been measured for binding to each of four different antibodies and to the ACE2 cell surface receptor [5, 6]. A mutation that displays reduced binding or loss of binding to any of the antibodies enables some amount of immune escape and thus is likely beneficial to the virus, whereas a mutation that displays reduced binding to the ACE2 cell surface receptor causes reduced viral fitness and is likely deleterious.

The 32,768 variants in the dataset where chosen because they represent all possible combinations of fifteen distinct mutations (2^15^ = 32, 768) that separate the Alpha and the Omicron variants of SARS-CoV-2 in the receptor binding domain of the viral spike protein [5, 6]. Thus, the dataset in effect maps out all possible evolutionary paths from Alpha to Omicron. All 32,768 variants were assessed for their binding affinity *K_D,app_* to class 1, 2, 3, and 4 antibodies (antibodies CB6, CoV555, REGN10987, and S309, respectively) and to the cell surface receptor ACE2.

We used the Hie *et al*. [10] model trained on the SARS-CoV-2 spike protein to calculate semantic change for all 32,768 variant sequences. We then correlated the semantic change values to the measured binding affinities as measured by *−* log *K_D,app_*. Larger values of *−* log *K_D,app_* indicate stronger binding, and a value below 6 indicates no detectable binding. For all four antibodies, we found a weak to moderate negative correlation between binding affinities above the limit of detection and semantic change (Fig. 5A). Thus, mutations with larger semantic change on average displayed weaker antibody binding. Moreover, for the three antibodies for which some mutations showed no binding at all (CB6, CoV555, and REGN10987), semantic change values where significantly larger on average for nonbinding variants than for binding variants (Fig. 5B). We note, however, that effect sizes where small in most cases. Correlation coefficients for three of the four antibodies fell between *−*0.12 to *−*0.25, 6.25% or less of the variance in binding explained by semantic change, and similarly semantic change for non-binding variants was on average only 0.175 units larger than for binding variants, while the standard deviation of semantic change values was between 0.23 and 0.28. Notably, for the S309 antibody, all assayed spike variants were active binders and demonstrated the strongest correlation between semantic change and binding affinity (Pearson’s *r* = *−*0.56, p-value *< .*0001, Fig. 5A).

**Figure 5:**
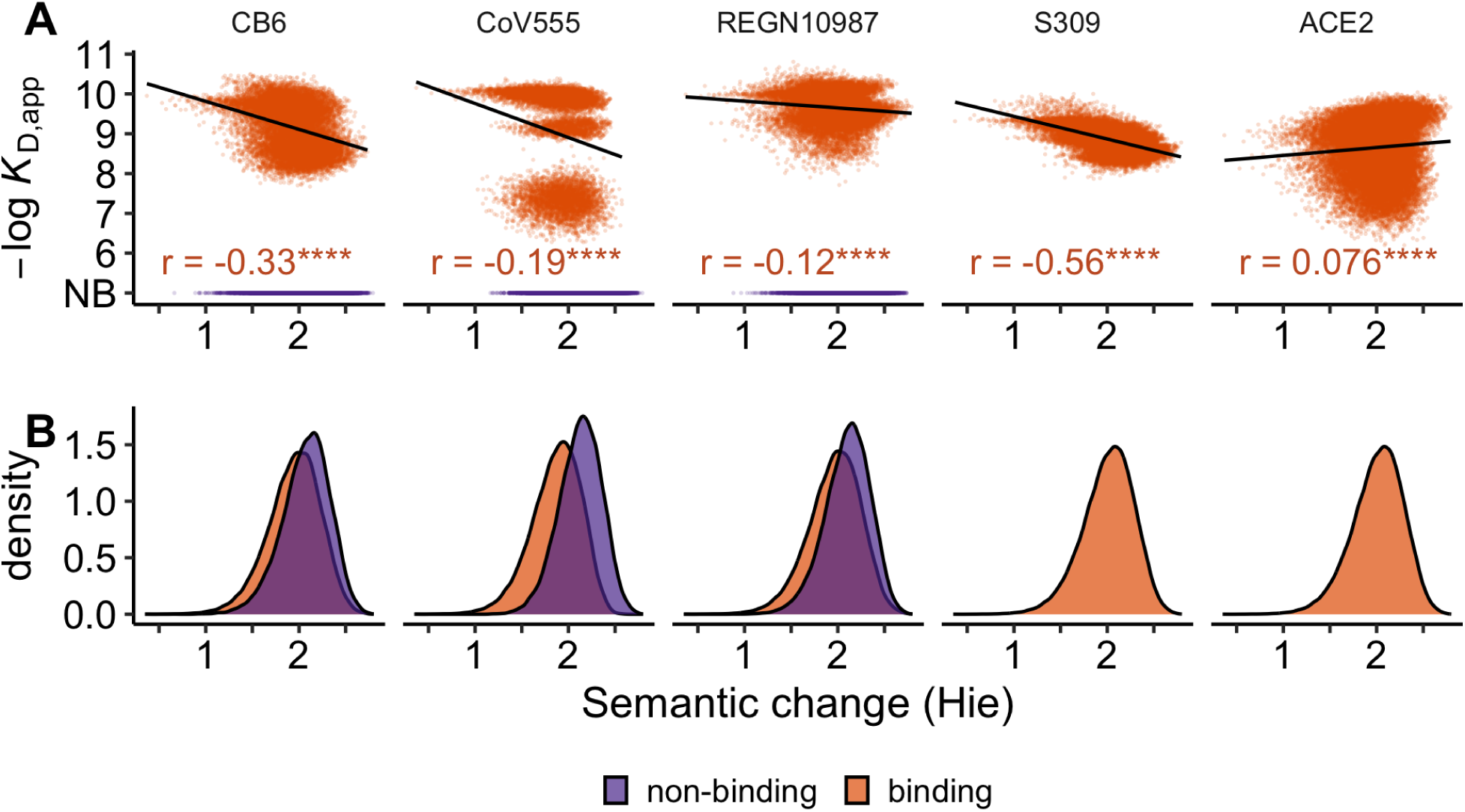
All possible combinations of 15 mutations defining the omicron BA.1 coronavirus spike protein had *K_D_* binding values measured for four antibodies [5] and ACE2 [6]. Semantic change was inferred from the Hie *et al*. [10] model. Colors represent mutations that confer escape (purple), or non-escape (orange), in the DMS experiment. (A) For each of spike’s binding partners, Pearson’s *r* was computed for all non-escape mutations since escape mutations were classified as being below the limit of detection for *K_D_*, so they are designated non-binding (NB). Significance is denoted with ****, indicating a p-value *<* 0.0001. (B) Density plots show the overlap of computed semantic change values between escape and non-escape mutations. No mutants failed to bind to antibody S309 and ACE2.

Surprisingly, the weakest correlation was between semantic change and the binding affinity to the ACE2 receptor (Pearson’s *r* = 0.076, p-value *< .*0001, Fig. 5A). This is notable in particular because the range of binding affinities to the ACE2 receptor among the 32,768 variants are comparable to the range of binding affinities to any of the antibodies. Thus, while the variants in this dataset clearly differed in their ability to bind to the ACE2 receptor, which impact viral entry, semantic change was unable to detect this variation.

We performed the same analysis on the combinatorial BA.1 mutations using the ESM2 model. Similar to the BiLSTM trained by Hie *et al*. [10], the correlation between binding affinity for three of the four antibodies (CB6, CoV555, and S309) was significantly different from zero, but was weak (Fig. S7A). For CB6, CoV555, and S309, ESM2 captured between roughly 6% and 21% of the variance in binding explained by semantic change (Pearson’s *r* between *−*0.26 and *−*0.46, p-value *<* 0.0001). Semantic change did not correlate with spike binding to REGN10987. While all variants bound to antibody S309, differences in means of semantic change between escape and non-escape mutations for the other three antibodies was 0.094 units on average, with standard deviations between 0.28 and 0.33, where all means of binders and non-binders fell within one standard deviation of one another (Fig. S7B). The ACE2 binding measures correlated weakly with semantic change from the ESM2 model (Pearson’s *r* = 0.079, p-value *<* 0.0001).

In summary, while semantic change was weakly correlated with loss of antibody binding, the effect size was rather small, and it did not simultaneously correlate with preserved binding to the ACE2 host receptor. Taken together, it would not be possible to reliably identify immune escape mutations based on their semantic change.

## 3 Discussion

We have systematically tested the language-of-viral-escape model by Hie *et al*. [10] using several new high-throughput datasets that have been made available since the original publication of the model and also using several additional models to calculate grammaticality scores. Overall, we have found that grammaticality is somewhat predictive of whether a mutation is viable or not, whereas semantic change is a less useful indicator of a mutation’s immune escape propensity. We have found that our results are broadly consistent across three different viral systems, SARS-CoV-2 spike, influenza HA, and HIV-1 env. We have also found that the bespoke models by Hie *et al*. [10] trained separately for each viral protein have not systematically outperformed generic, pretrained sequence-or structure-based foundation models. Finally, we have found that for the task of predicting mutant viability (i.e. grammaticality), structure-based models seem to outperform pLMs, and their performance can be further improved when fine-tuned on experimental ΔΔ*G* datasets.

Importantly, we have seen a major difference in performance between grammaticality scores calculated using the masked or the unmasked ESM2 model. While scores obtained from the masked model always showed reduced grammaticality for nonviable mutations, and in the case of influenza HA were competitive with the structure-based models in terms of the magnitude of predicted difference between viable and nonviable mutations, results were inconclusive or pointed the opposite direction for the unmasked model. We believe the masked approach is the appropriate one and should be used, and we discourage the use of the unmasked approach. ESM models, which are built on a BERT-style [32] encoder-only transformer architecture, utilize masked language modeling (MLM) as the training objective [13]. MLM is a fundamental self-supervised learning technique that enables learning the identity of the masked positions from the sequence context—in proteins, this is the dependencies between amino acids. When we ask the model to make a prediction for the masked site, it predicts all amino-acid propensities at once and conditional on each other. By contrast, in the unmasked approach, the model is asked to make multiple separate predictions, one for each possible sequence variant, and these predictions are not conditional on each other. And more importantly, the training procedure of ESM models for unmasked sites has biased the model towards returning either the input token or any one of the potential output tokens chosen from a uniform distribution [13]. Therefore, there is no good *a priori* reason why the unmasked inference procedure should be successful, and our analysis here has shown that it is not.

It is notable that zero-shot predictions from structure-based models have systematically performed better than sequence-based models in differentiating viable from nonviable mutations, including bespoke models. In a prior study, we had compared pretrained sequence-and structure-based models for their ability to predict the wild-type residue when masked [23]. In that study, we had found that both model types displayed roughly comparable performance, even though we saw some differences in the specific amino acids for which they made more or less reliable predictions. The discrepancies in these two different studies highlight that two model types can have comparable performance on one task yet differing performance on another. To correctly predict the masked wild-type amino acid at a particular residue, a model needs to assign a high probability to the wild-type amino acid and low probabilities to all other amino acids. However, the specific probabilities assigned to the other amino acids do not matter as long as they are lower than the probability of the wild-type amino acid. By contrast, to correctly predict nonviable mutations at a residue, a model needs to consistently assign low probabilities to the nonviable amino acids and higher probabilities to the viable amino acids. Our results here suggest that models taking structure as input appear to systematically perform better at this task. This is further supported by our previous studies demonstrating their ability to successfully identify viable mutations and guide protein engineering campaigns for a variety of proteins [21, 33–37].

We have also found that across all models and viral systems, the Stability Oracle [30], which predicts ΔΔ*G* values, has most consistently applied low scores to nonviable mutations and high scores to viable mutations. This observation touches upon the biophysical meaning of grammaticality. The original vision of the language-of-viral-escape model [10] was that language models could implicitly learn which mutations are viable and which are not and encode this information in their predicted amino acid propensities. What we have seen here is that even though language models do have this capability to some extent, structure-based models are more capable, and explicit fine-tuning of a structure model to predict the biophysical quantity ΔΔ*G* seems to be the most effective approach. This observation is consistent with the long-standing observation that destabilizing mutations are the primary culprit for loss of function in proteins [9, 38–42]. Therefore, for the purpose of predicting which mutations are viable and which are not, we would recommend to use a good ΔΔ*G* predictor instead of a language model and simply categorize the destabilizing mutations as likely nonviable. Here, we have only considered Stability Oracle, as it is one of the best deep learning frameworks for ΔΔ*G* prediction currently available, but we expect that other predictors, such as FoldX [43], PoPMuSiC [44], or Rosetta ddG [45], would perform similarly, and in proportion to their ability to predict accurate ΔΔ*G* values.

In contrast to grammaticality, we have found that semantic change does not seem to perform as originally expected. It does not consistently differentiate escape from non-escape mutations, and correlations between binding constants and semantic change are weak even if significant. And even absent these results, the concept of semantic change suffers from a fundamental problem [46]: If a large semantic change coincided with loss of binding to an antibody, why shouldn’t it also coincide with loss of binding to the host receptor? In both cases, the surface of the spike protein has been altered sufficiently that binding is no longer possible. Instead, we have found here that it does not strongly correlate with either. While distances in embedding space have been useful in some applications, such as inferring GO Terms [47], it appears that the complex phenotype of immune escape is not sufficiently represented by just distance in embedding space (i.e., semantic change). Instead, strategies that have worked well in a variety of different applications consist of transferring or fine-tuning the hidden representations of a language model to predict the phenotype of interest [30, 48–51]. An additional strategy includes learning to extract a phenotype-aware representation from the initial hidden representation that is enriched for the specific downstream application [52]. Alternatively, or in combination with such approaches, one can also build biophysical models of protein folding and binding and calibrate with experimental measurements of binding constants or measurements of viral fitness [9, 53].

One recent study [11] explored a variation of the language-of-viral-escape model to assess whether variants of concern (VOCs) are distinct from non-VOCs in SARS-CoV-2. Notably, they did not use the bespoke models of Hie *et al.* [10] and only computed grammaticality and semantic change from ESM2 embeddings. Their main focus was the change of these quantities over evolutionary time, but they also assessed correlations of semantic change and grammaticality with viral escape. Their results were broadly consistent with our findings here: Correlations are significant but weak. Moreover, they did not use a masking approach to calculate grammaticality, and we believe this may explain why they also observed weak correlations between grammaticality and physical measures of protein viability such as ΔΔ*G*.

In summary, we have found that the language-of-viral-escape framework [10], as currently developed, is not sufficient to accurately predict immune escape mutations. While the concept of grammaticality is informative about mutant viability, the concept of semantic change provides little information about whether or not a mutant will likely confer immune escape. Moreover, even for mutant viability, long-standing quantities such as stability changes appear to be more useful than grammaticality scores extracted from language models. We believe that the way forward for AI applications in viral evolution is to fine-tune (in a supervised framework) representations from pretrained models to predict specific phenotypes rather than rely on zero-shot predictions extracted from pretrained models only.

## 4 Methods

### 4.1 Language models

Semantic change is defined as the distance between embeddings of the wild-type and mutant sequences. We define this in the same way of Hie *et al.* [10]:

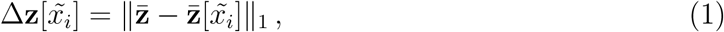

where **z** is an *L×d* embedding for a wild-type sequence of length *L* and **z**[*x̃_i_*] is the embedding of a sequence mutated to token *x* at locus *i*. The mean embeddings, **z̅**, are computed across sites resulting in 1 *× d* vectors. After taking the difference of these two vectors, the *l*^1^ norm is then computed (i.e. the sum of the absolute values of distances from **z̅** to **z̅**[*x̃_i_*]).

Grammaticality in the LLM context is defined as the emitted probability from the protein language model for the input sequence,

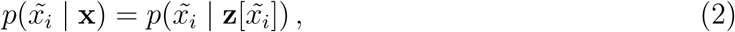

where *p*(*x̃_i_* | **x**) is the probability of observing mutation *x̃* at locus *i* from the amino acid alphabet **x**. When considering a mutated sequence, one can either use a masked or unmasked inference approach. Hie *et al*. [10] computed the probability over the sequence using their trained BiLSTM without masking, and in doing so extracted the probability of the mutant token(s). Their approach is aware of the identity of every token in the sequence when inferring the probability from the embedding, so it is unmasked. In addition to using the embeddings from their bespoke-trained model, we used the embeddings from ESM2 to obtain unmasked grammaticality predictions. For ESM2, we also implemented a masked grammaticality by taking a mutated sequence and swapping the amino acid token of the mutated locus with the <mask> token. The model then must compute the probability of observing each amino acid at the masked locus using only the embeddings of the remaining unmasked amino acids. These probabilities serve as the grammaticality scores for each amino acid at each site.

Semantic change and grammaticality measures were inferred from multiple LLMs. We used the published outputs for influenza and HIV [10], but since SARS-CoV-2 used a different reference sequence in the DMS [7], we used the bespoke pre-trained coronavirus model [10] to produce novel results. We then used ESM2 (esm2 t30 150M UR50D) [13] with the same model architecture to obtain semantic change and grammaticality scores from this generally trained model. Additionally, we wrote Python scripts to allow for masking of tokens when inferring grammaticality from the ESM2 model using the transformers Python library. Scripts to perform inference of semantic change and grammaticality are available at https://github.com/allmanbrent/NLP_viral_escape/tree/main/language_models.

### 4.2 AlphaFold

For the structure-based models, we needed protein structures as inputs. For all three viruses, we inferred structures using AlphaFold2. As all three viruses exist as homotrimer in their natural state, we used AlphaFold2’s multimer functionality [54, 55], as implemented in the ColabFold notebook: https://colab.research.google.com/github/sokrypton/ ColabFold/blob/main/AlphaFold2.ipynb. The structure files are available on https://github.com/allmanbrent/NLP_viral_escape/tree/main/data.

### 4.3 MutComputeX and MutRank

MutComputeX [21] is a self-supervised 3-dimensional residual neural network (3DResNet) trained on *∼*2.1 millions protein masked microenvironments sampled from *∼*23K protein structures. The 3DResNet is trained with the cross entropy loss to predict the masked amino acid in the center of the masked microenvironment. The trained model outputs likelihood of each amino acid for a particular microenvironment. We used these likelihoods as our grammaticality scores to compare against other models. The inference pipeline can be found here: https://github.com/danny305/MutComputeX/blob/master/scripts/generate_predictions.py.

MutRank [22] is a self-supervised 3-dimensional graph transformer that adds a second training step to a graph-transformer analog of MutComputeX. In the second step, it trains a regression head that takes a “FromAA” and “ToAA” CLS tokens as additional inputs to the masked microenvironment using the EvoRank loss on the MutComputeX training set to predict the rank score between the “FromAA” and “ToAA”. During inference, for each masked microenvironment we set the “FromAA” CLS token to the wildtype amino acid and predict the rank score for all 20 amino acids by setting them as the “ToAA” CLS token. We used the 20 rank scores obtained for each microenvironment as our grammaticality scores to compare against other models. The inference pipeline can be found here: https://github.com/danny305/StabilityOracle/blob/master/scripts/run_stability_oracle.py

### 4.4 Stability Oracle

Stability Oracle [30] is a structure-based graph-transformer model that supervise fine-tunes a graph transformer analog of MutComputeX on empirical ΔΔ*G* values of the cDNA117K dataset. The architecture is identical to the MutRank architecture but rather than training the regression head with the EvoRank loss on the MutComputeX training set, the regression head is trained (and the graph-transformer backbone is fine-tuned) with the huber loss on the cDNA117K dataset. During inference, for each masked microenvironment we set the “FromAA” CLS token to the wildtype amino acid and predict ΔΔ*G* values for all 20 amino acids by setting them as the “ToAA” CLS token. We used the 20 ΔΔ*G* values obtained for each microenvironment as our grammaticality scores to compare against other models. The values are reversed because positive ΔΔ*G* values correspond to more destabilizing mutations (lower grammaticality) and negative ΔΔ*G* values correspond to more stabilizing mutations (higher grammaticality). The inference pipeline can be found here: https://github.com/danny305/StabilityOracle/blob/master/scripts/run_stability_oracle.py

### 4.5 Statistical testing

All statistical tests were performed in R. Scripts to modify the data for boxplots and perform Mann-Whitney U test are available on Github. For all figures, model names have been abbreviated and here we briefly describe what each label means. “Grammaticality (Hie)” refers to the probabilities emitted by the LLM used in Hie *et al*. [10] that was trained on coronavirus sequences. “Grammaticality (ESM2, unmasked)” refers to the probabilities emitted by the LLM scheme used by Hie *et al*. [10] with the general ESM2 model. The model is unmasked because the protein sequence tokens were not masked when the model determines the probability of a token belonging in a particular locus. “Grammaticality (ESM2, masked)” refers to the probabilities emitted by ESM2, but the sequence tokens are masked. “MutComputeX” refers to the amino acid probabilities emitted by the structure-based model from d’Oelsnitz *et al*. [21]. “Stability Oracle” refers to the ΔΔ*G* values predicted by Diaz *et al*. [30]. Note that the ranks for Stability Oracle are reversed since more negative ΔΔ*G* values correspond to higher stability and thus higher grammaticality. “Semantic change (Hie)” refers to the normalized differences between wild-type and mutant sequence embeddings emitted by the coronavirus LLM used in Hie *et al*. [10]. “Semantic change (ESM2)” refers to the normalized differences between wild-type and mutant sequence embeddings emitted by the general ESM2 LLM.

### 4.6 Coronavirus spike glycoprotein

The spike sequence used as the template for computationally mutating every residue is available on NCBI with GenBank ID QHD43416.1. The Wuhan spike sequence used for the 15 combinatorial Omicron mutations is available on NCBI with GenBank ID YP 009724390.1. The library of mutants used as input to the language models, the template sequences, and the homotrimer protein structure used as input to MutCompute, MutRank, and Stability Oracle are available at https://github.com/allmanbrent/NLP_viral_escape/tree/main/data/cov.

We utilized data from a DMS experiment of the SARS-CoV-2 spike protein where a pseudovirus system was used to determine the impact of mutations on antigenic escape [7]. In this experiment, a subset of sites was tested for their antigenic escape phenotype and some mutations did not confer a viable protein. These were denoted by a lack of escape measurement from the experiment. So here, viable mutations were classified as either escape or non-escape by their reported escape fraction. Those with an escape fraction above 0.5 were designated escape. These data are available at https://github.com/allmanbrent/NLP_viral_escape/tree/main/data/cov/starr_dms.

We also used data from previously published DMS experiments where the 15 mutations that define the Omicron SARS-CoV-2 strain were tested for binding affinity for four antibodies [5] and the cell surface receptor ACE2 [6]. These data are available at https://github.com/allmanbrent/NLP_viral_escape/tree/main/data/cov/omicron_experiments

### 4.7 Influenza A hemagglutinin protein

The HA sequence used as the template for computationally mutating every residue is identical to the NCBI sequence with GenBank ID QDQ43389.1. The library of mutants used as input to the language models, the template sequence, and the homotrimer protein structure used as input to MutCompute, MutRank, and Stability Oracle are available at https://github.com/allmanbrent/NLP_viral_escape/tree/main/data/flu. Note that we use the previously published results from Hie *et al*. [10] for the BiLSTM language model.

We used previously published DMS of the H1 hemagglutinin protein of influenza A/WSN/1933 [4]. They determined the mutational tolerance of each site along the entire protein sequence. The results of this experiment were amino acid preferences at each site (excluding the start codon) for all amino acids; i.e. the expected post-selection frequency of all 20 amino acids at each site for all possible single mutant sequences. From these data, we classified mutations as resulting in viable or not viable proteins. The data we used from this experiment are available at https://github.com/allmanbrent/NLP_viral_escape/tree/main/data/flu/escape_doud2018.

We defined viable mutations as those having an amino acid preference above 0.001. To make this determination, we looked at the curve representing the ranked amino acid preferences and note the behavior changes at 0.1 and 0.001 (Fig. S8). Of the 1222 mutations with an amino acid preference above 0.1, 482 are wild-type (Fig. S8). This leaves 82 wild-type mutations classified as not viable if we chose a cutoff of 0.1. The experimental per-codon sequencing error rate is somewhere between 0.0002 and 0.0005, and the observed nonsynonymous post-selection frequency is approximately 0.0008 [4]. Thus, we chose the cutoff of 0.001 to be a slightly more stringent classifier of viability than the bounds for sequencing error. The data we use from this experiment are available at https://github.com/allmanbrent/NLP_viral_escape/tree/main/data/flu/fitness_doud2016. The code used to define the viability of mutations is available at https://github.com/allmanbrent/NLP_viral_escape. Escape fractions were obtained in the DMS of A/WSN/1993 [17], but a simple nu-merical cutoff is insufficient to define escape mutations in this case. We used dms tools2 [56] to classify mutations as escape or non-escape for each of the antibodies tested. We then looked across antibodies and if a variant confers escape under any one antibody selection scheme, then it is considered an escape mutation. The data we used from this experiment are available at https://github.com/allmanbrent/NLP_viral_escape/tree/main/data/flu/escape_doud2018. The code used to define escape mutations is available at https://github.com/allmanbrent/NLP_viral_escape.

### 4.8 HIV-1 envelope glycoprotein

We used the BG505.W6M.C2.T332N strain of env which has DMS experiments testing viability [19] and antigenic escape [18]. The library of mutants used as input to the language models, the template sequence, and the homotrimer protein structure used as input to MutCompute, MutRank, and Stability Oracle are available at https://github.com/allmanbrent/NLP_viral_escape/tree/main/data/hiv. Note that we used the previously published results from Hie *et al*. [10] for the BiLSTM language model.

Similar to the influenza experiment described above, 670 sites from the HIV BG505 strain env protein have been previously mutagenized and then their amino acid preferences were measured based on observed frequencies from deep sequencing taken from *in vitro* cell passage [19]. To define viability from these amino acid preferences, we again looked at the rank of the preferences (Fig. S9). Like influenza, the inflection points for the behavior of these ranked data shift at amino acid preferences of 0.1 and 0.001. With a cutoff of 0.1, 243 of the wild-type mutations are classified as not viable. Therefore, we used the cutoff of 0.001 since this captures all wild-type mutations and is slightly more stringent than proposed error rates [19]. The data we use from this experiment are available at https://github.com/allmanbrent/NLP_viral_escape/tree/main/data/hiv/Haddox_supp. The code used to define the viability of mutations is available at https://github.com/allmanbrent/NLP_viral_escape.

We used escape fractions from previously published DMS on BG505 HIV where the virus underwent antibody selection [18]. We used dms tools2 [56] to classify mutations as escape or non-escape for each of the antibodies tested. We then look across antibod-ies and if a variant confers escape under any one antibody selection scheme, then it is considered an escape mutation for our purposes. The data we used from this experiment are available at https://github.com/allmanbrent/NLP_viral_escape/tree/main/data/hiv/Dingens_ab_escape. The code we used to define escape mutations is available at https://github.com/allmanbrent/NLP_viral_escape.

## Data Availability

The code used to reproduce the analysis shown in this paper is available at https://github.com/allmanbrent/NLP_viral_escape. The viral sequences and structures used as input can also be found in the same Github repository.

## Acknowledgments

We would like to thank Anastasiya Kulikova for helpful conversations and assistance with the LLM computing. We thank the Institute for Foundational of Machine Learning (IFML), the Texas Advanced Computing Center, and the Biomedical Research Computing Facility at the University of Texas at Austin for the computing resources used to do the analyses in this manuscript. We would like to thank AMD for the donation of critical hardware and support resources from its HPC Fund.

## Funding

This study was supported by NSF award DEB 2200169. C.O.W. was also supported by the Jane and Roland Blumberg Centennial Professorship in Molecular Evolution and the Dwight W. and Blanche Faye Reeder Centennial Fellowship in Systematic and Evolutionary Biology at The University of Texas at Austin. D.J.D was supported by the NSF AI Institute for Foundations of Machine Learning (IFML).

## Conflicts of Interest

D.J.D. has a financial relationship with Intelligent Proteins LLC, which uses AI models for protein engineering.

B.A., L.V., and C.O.W. declare no competing interests.

## Supplementary Materials

Supplementary Materials are at the end of this document.

## 5. Supplementary Material

**Figure S1:**
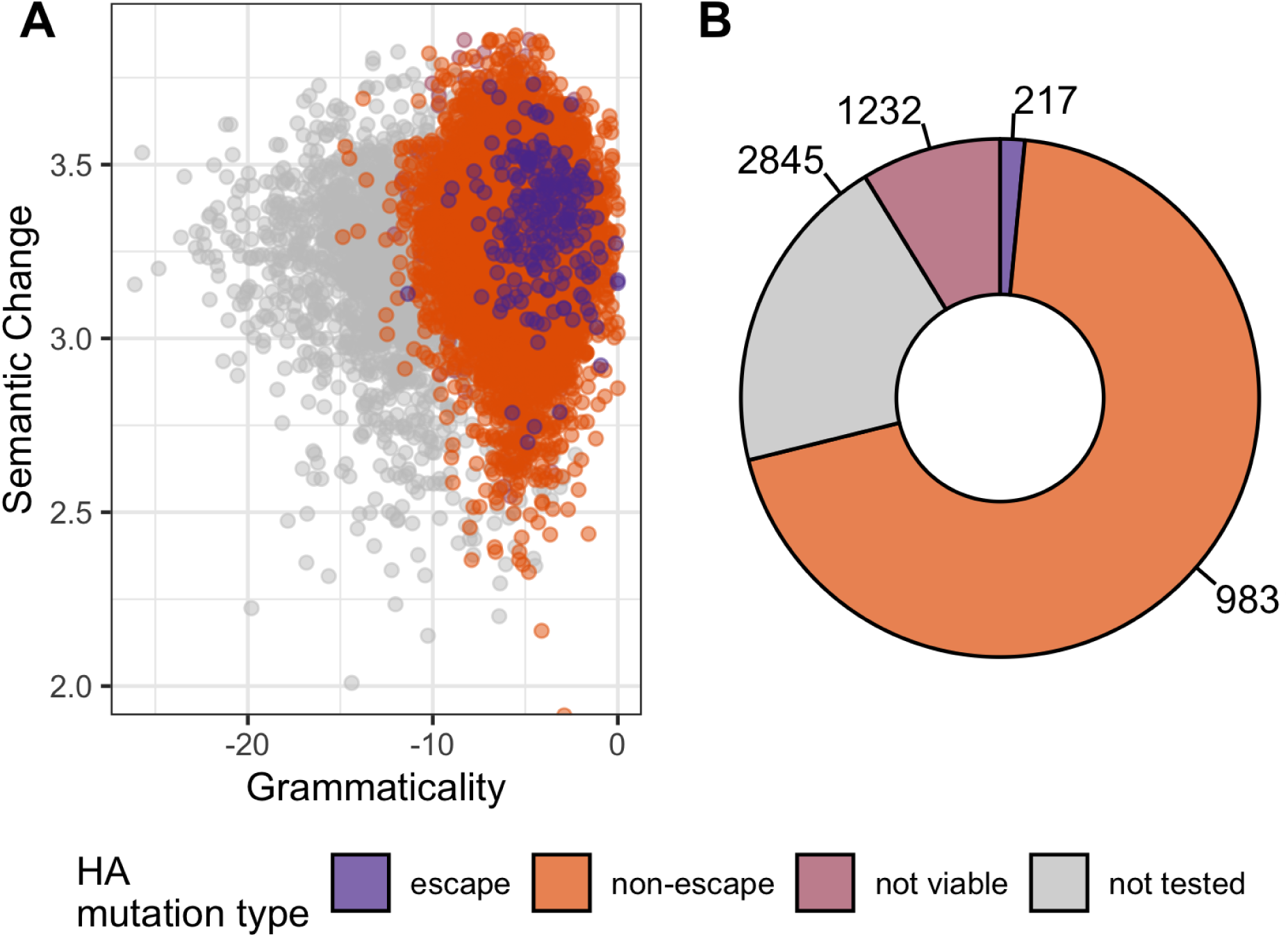
Semantic change and grammaticality scores of mutations predicted by Hie *et al*. [10] attempt to provide insight into antigenic escape. Deep mutational scanning data [4, 17] validate propensity for a mutation to escape antibody pressure, but also give insight into the viability of mutations. Purple represents mutations classified as escape. Orange represents mutations that did not successfully escape antibody pressure. Pink represents mutations that render the virus not viable. Gray represents mutations that were not tested for antigenic escape.

**Figure S2:**
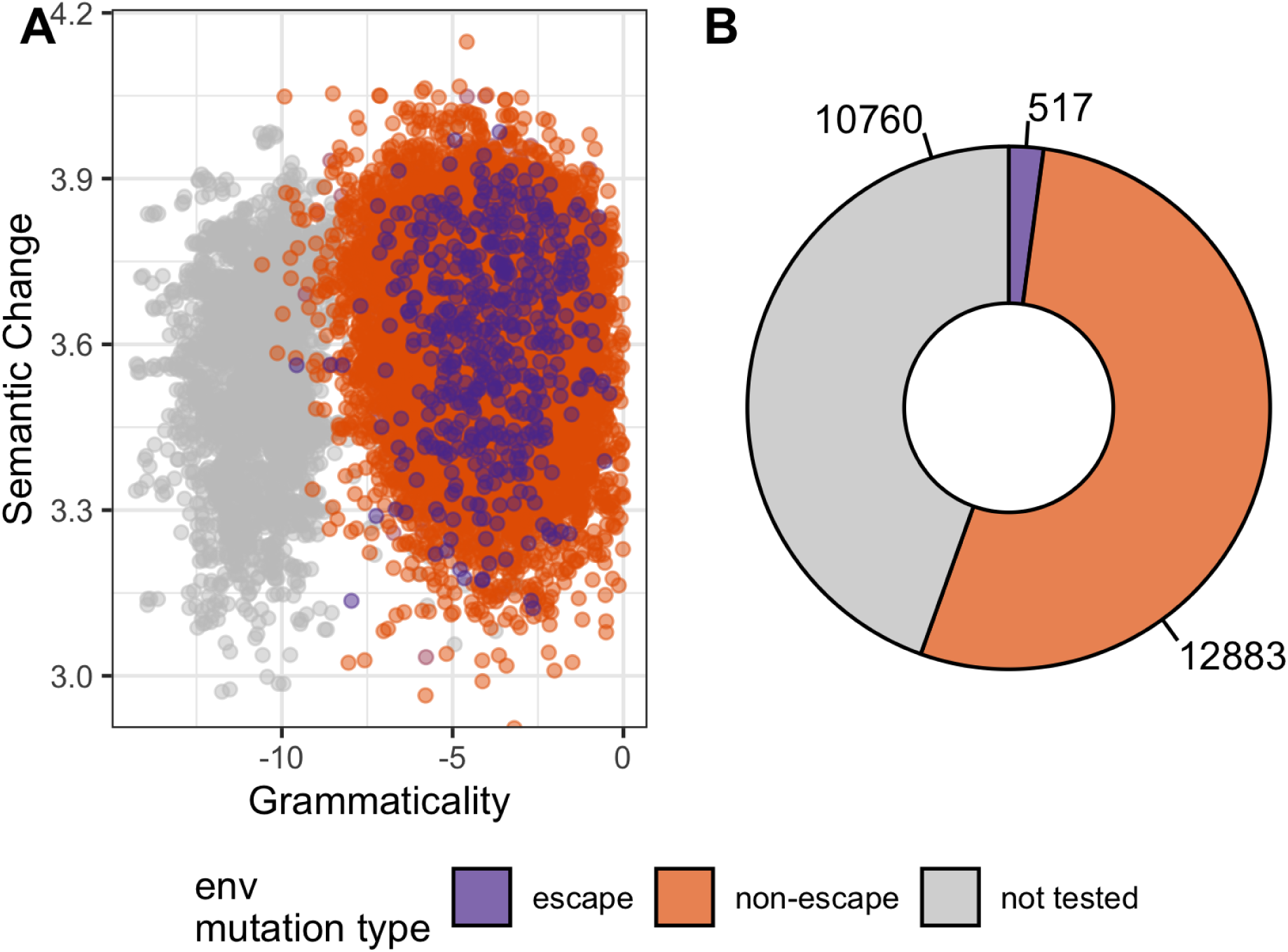
Semantic change and grammaticality scores of mutations predicted by Hie *et al*. [10] attempt to provide insight into antigenic escape. Deep mutational scanning data [18] validate propensity for a mutation to escape antibody pressure, but also give insight into the viability of mutations. Purple represents mutations classified as escape. Orange represents mutations that did not successfully escape antibody pressure. Gray represents mutations that were not tested for antigenic escape.

**Figure S3:**
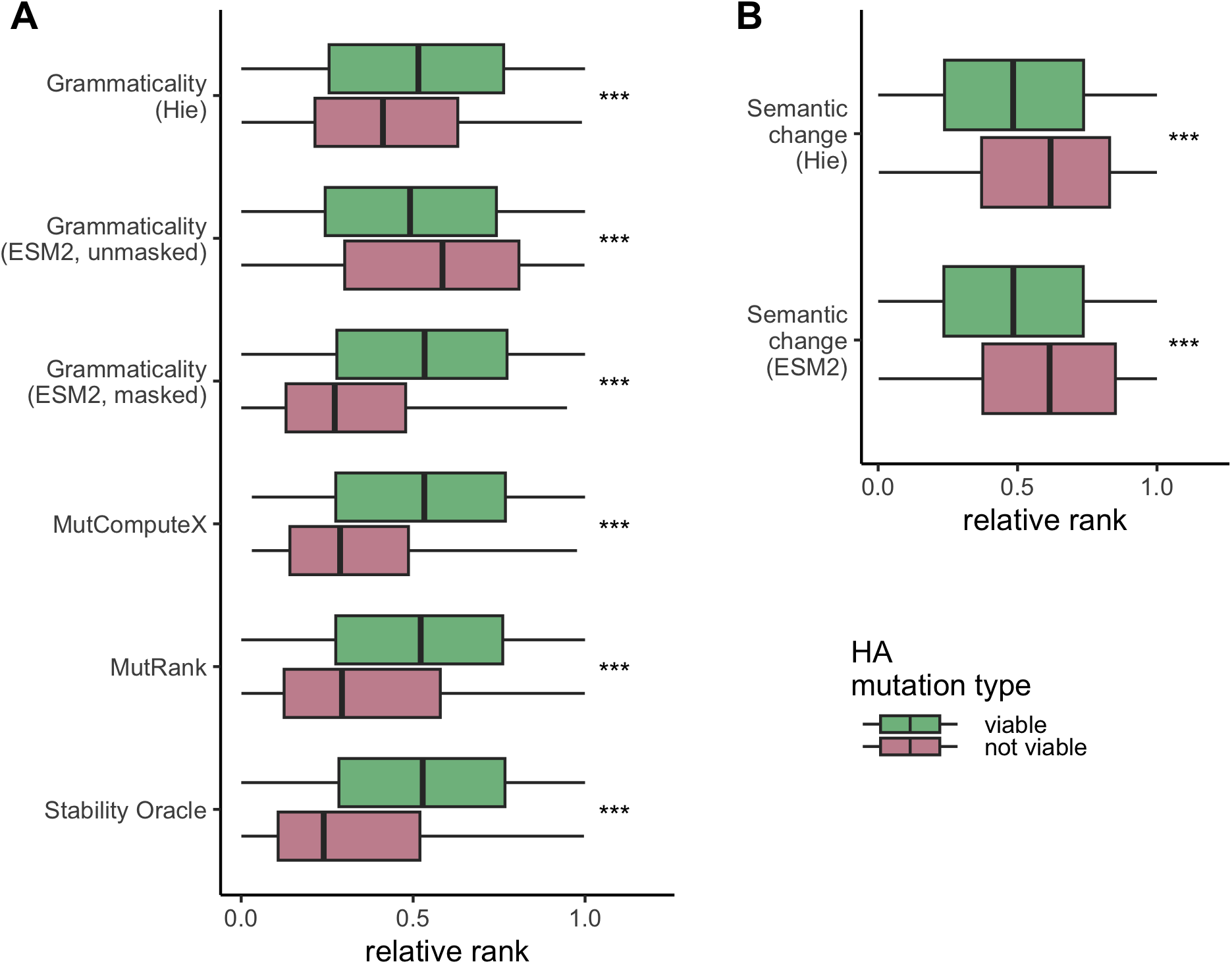
All possible mutations of the influenza A virus hemagglutinin protein DMS experiment [4] tested under different models. Colors represent mutations that are viable (green) or not viable (pink) in the DMS experiment. The values predicted for each mutation are ranked and then normalized to be between 0 and 1. (A) Grammaticality scores for each of the six models. Note that the ranks for Stability Oracle are reversed since small ΔΔ*G* values are consistent with higher stability. (B) Semantic change scores for both the Hie *et al*. [10] model and the ESM2 model. Results of the Mann-Whitney U rank test are indicated as: NS *→* not significant, *∗ → p < .*05, *∗ ∗ → p < .*01, *∗ ∗ ∗ → p < .*001.

**Figure S4:**
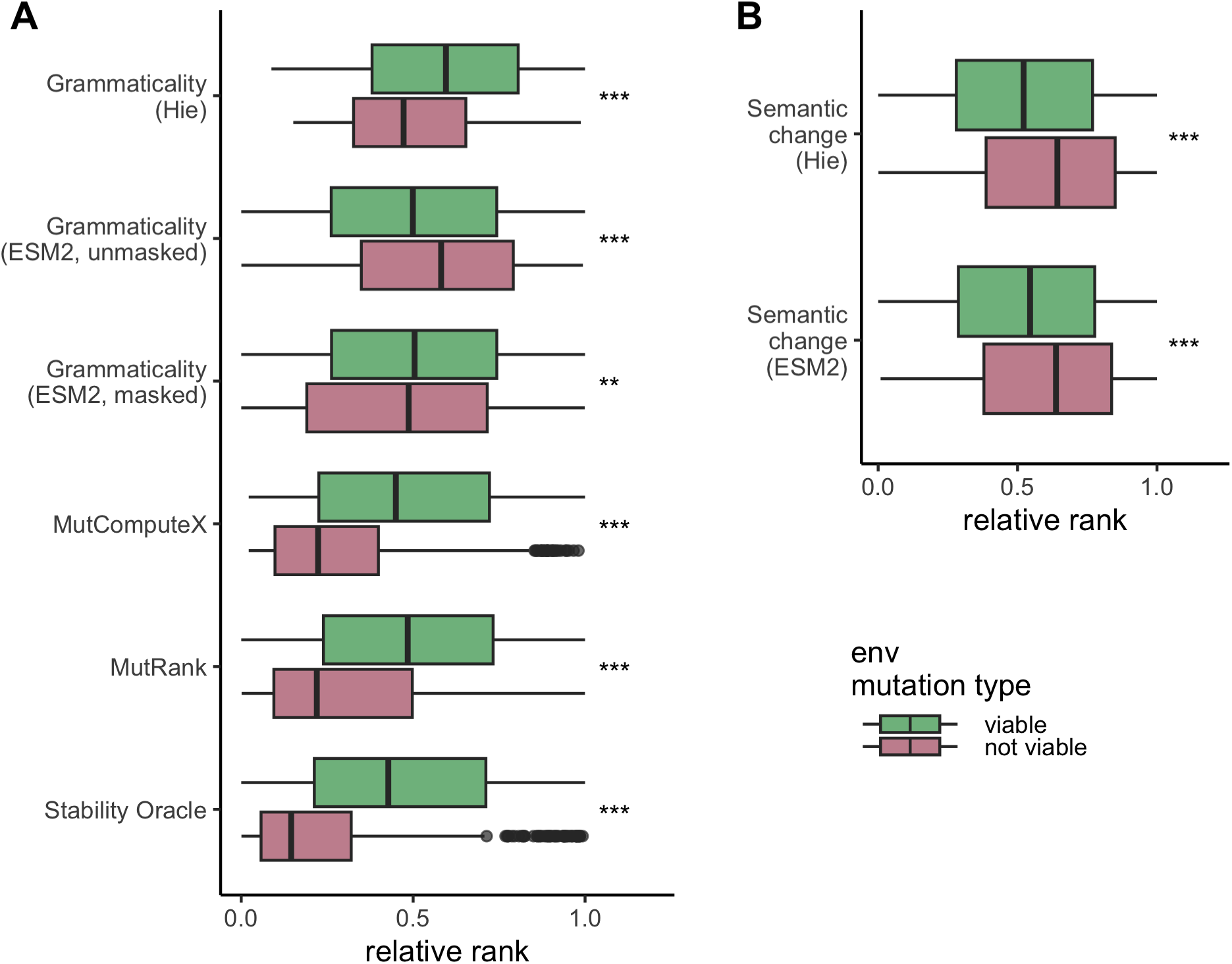
All possible mutations of the HIV envelope glycoprotein DMS experiment [19] tested under different models. Colors represent mutations that are viable (green) or not viable (pink) in the DMS experiment. The values predicted for each mutation are ranked and then normalized to be between 0 and 1. (A) Grammaticality scores for each of the six models. Note that the ranks for Stability Oracle are reversed since small ΔΔ*G* values are consistent with higher stability. (B) Semantic change scores for both the Hie *et al*. [10] model and the ESM2 model. Results of the Mann-Whitney U rank test are indicated as: NS *→* not significant, *∗ → p < .*05, *∗ ∗ → p < .*01, *∗ ∗ ∗ → p < .*001.

**Figure S5:**
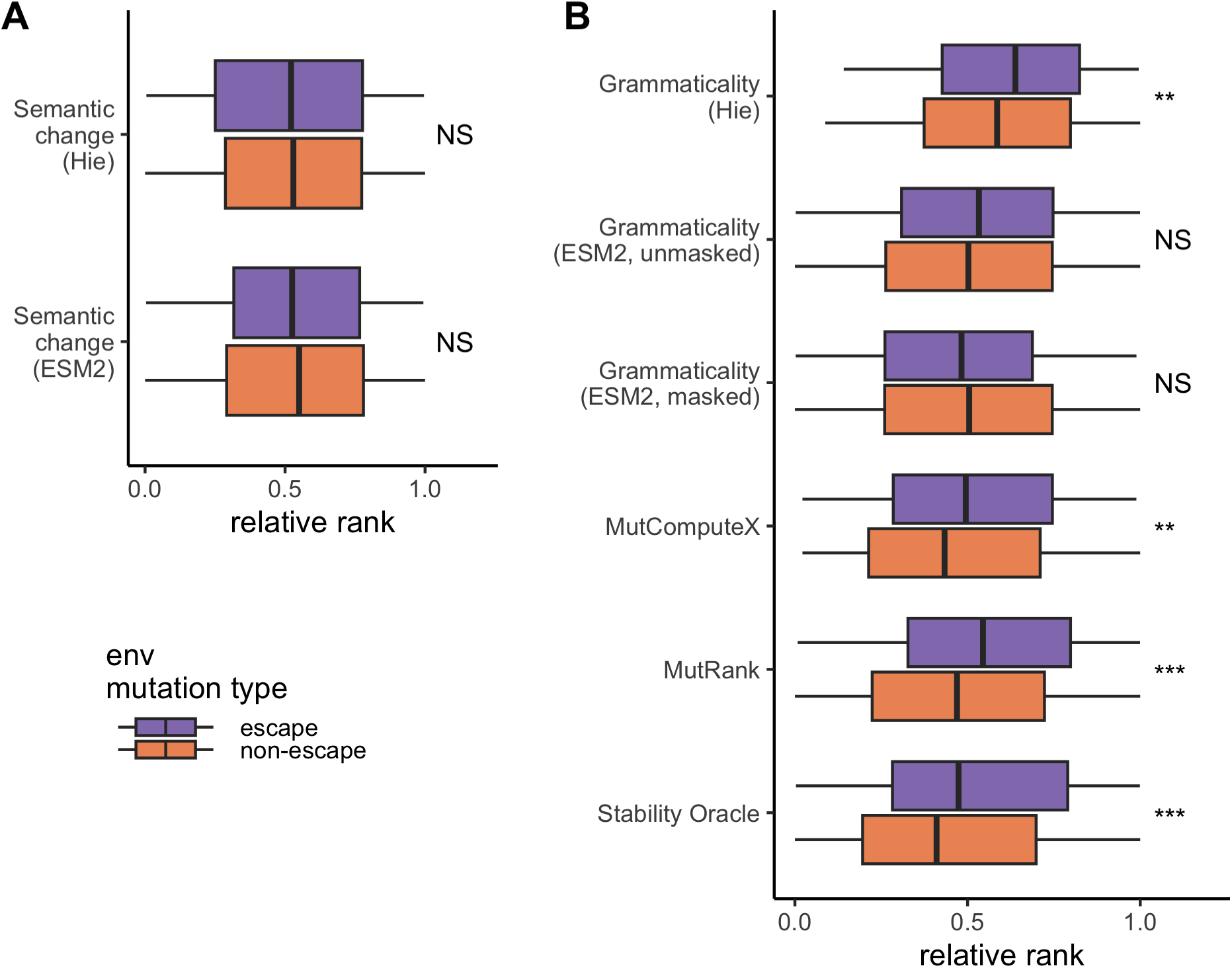
All possible mutations of the HIV envelope glycoprotein DMS experiment [18] tested under different models. Colors represent mutations that confer escape (purple) or non-escape (orange) in the DMS experiment. The values predicted for each mutation are ranked and then normalized to be between 0 and 1. (A) Semantic change scores for both the Hie *et al*. [10] model and the ESM2 model. (B) Grammaticality scores for each of the six models. Note that the ranks for Stability Oracle are reversed since small ΔΔ*G* values are consistent with higher stability. Results of the Mann-Whitney U rank test are indicated as: NS *→* not significant, *∗ → p < .*05, *∗ ∗ → p < .*01, *∗ ∗ ∗ → p < .*001.

**Figure S6:**
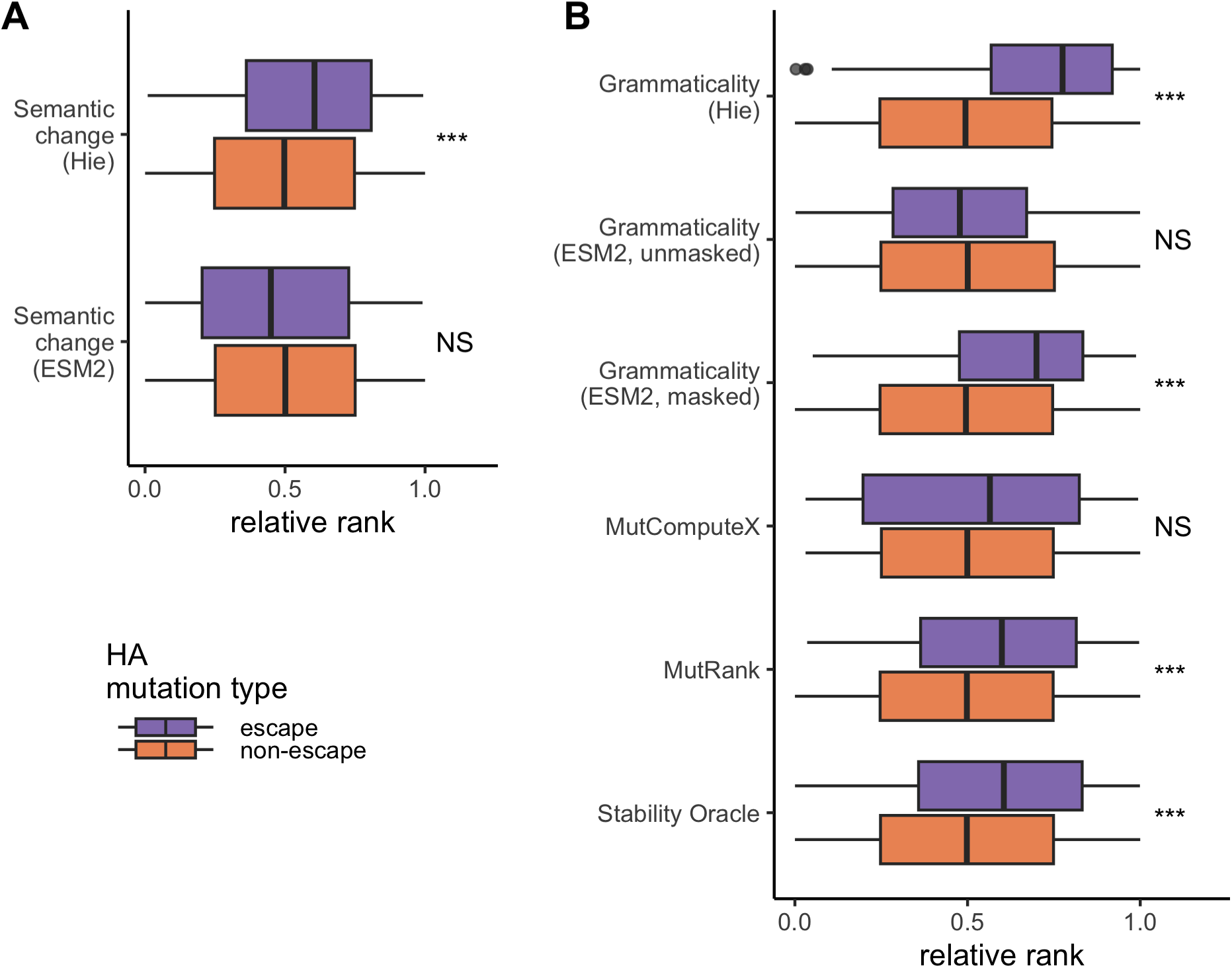
All possible mutations of the influenza A virus hemagglutinin protein DMS experiment [17] tested under different models. Colors represent mutations that confer escape (purple) or non-escape (orange) in the DMS experiment. The values predicted for each mutation are ranked and then normalized to be between 0 and 1. (A) Semantic change scores for both the Hie *et al*. [10] model and the ESM2 model. (B) Grammaticality scores for each of the six models. Note that the ranks for Stability Oracle are reversed since small ΔΔ*G* values are consistent with higher stability. Results of the Mann-Whitney U rank test are indicated as: NS *→* not significant, *∗ → p < .*05, *∗ ∗ → p < .*01, *∗ ∗ ∗ → p < .*001.

**Figure S7:**
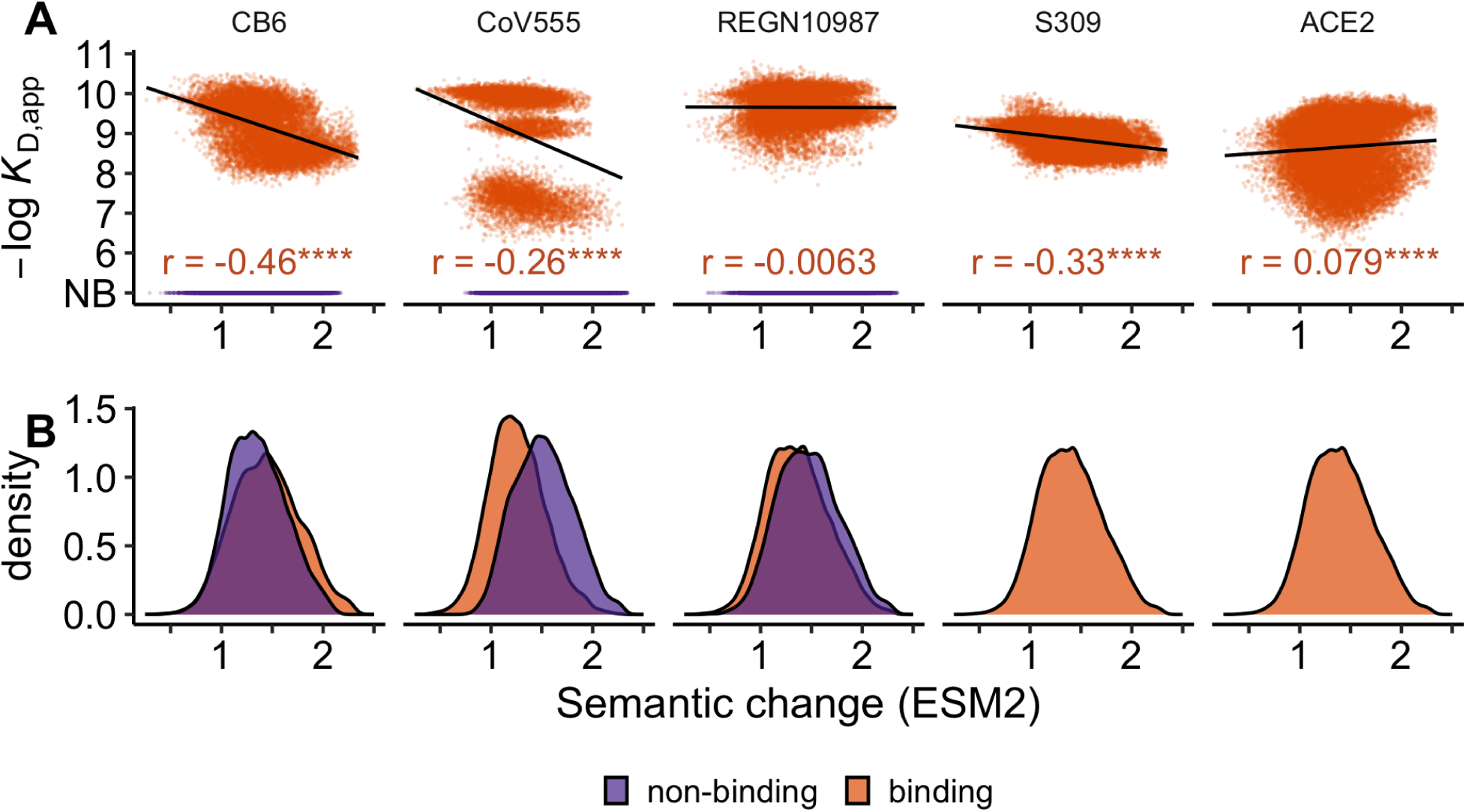
All possible combinations of 15 mutations defining the omicron BA.1 coronavirus spike protein had *K_D_*binding values measured for four antibodies [5] and ACE2 [6]. Semantic change was inferred from ESM2 [13] is presented on a *log*_10_ scale. Colors represent mutations that confer escape (purple), or non-escape (orange), in the DMS experiment. (A) For each of spike’s binding partners, Pearson’s *r* was computed for all non-escape mutations since escape mutations were classified as being below the limit of detection for *K_D_*, so they are designated non-binding (NB). Significance is denoted with ****, indicating p-value *<* 0.0001. (B) Density plots show the overlap of computed semantic change values between escape and non-escape mutations. No mutants failed to bind to antibody S309 and ACE2.

**Figure S8:**
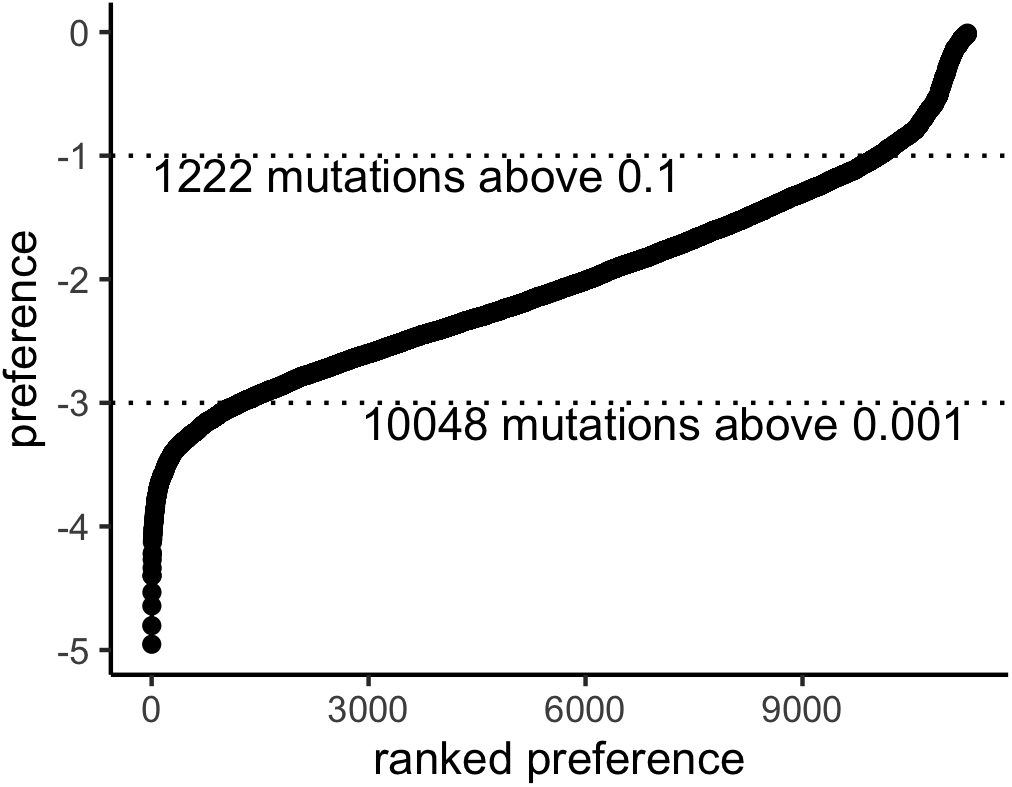
Amino acid preferences of all mutations from A/WSN/1993 with the exception of the start codon, ranked [4]. Horizontal dotted lines represent putative cutoffs for defining viable mutations.

**Figure S9:**
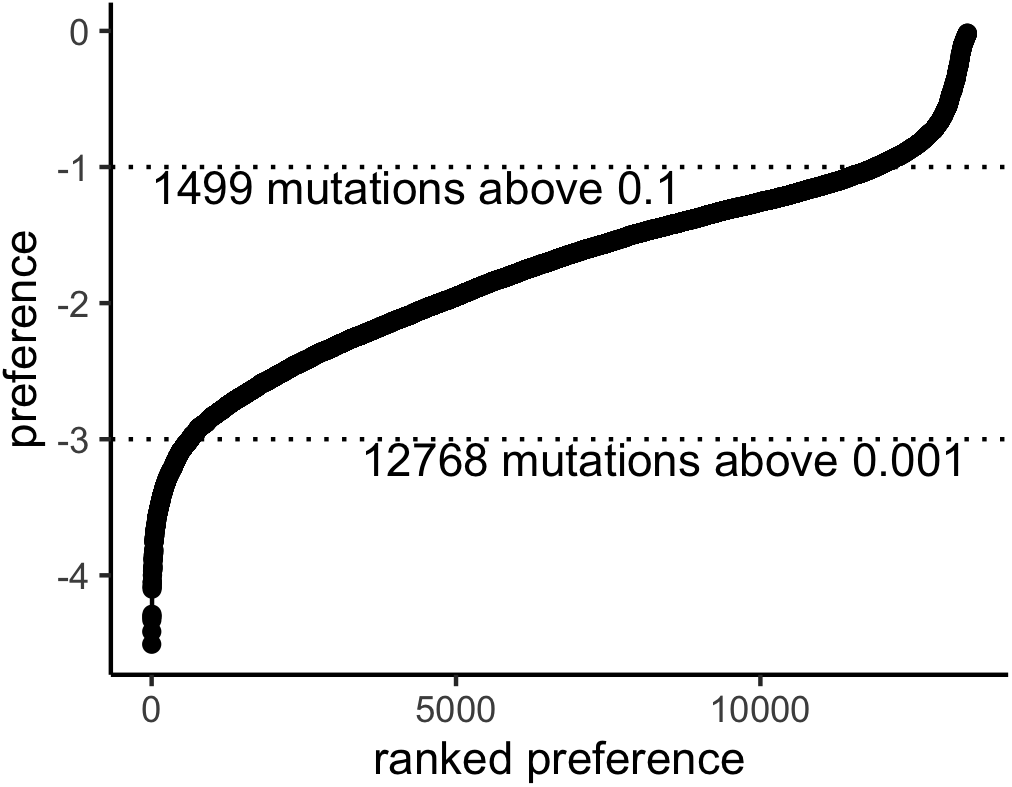
Amino acid preferences of mutations 670 sites BG505 env with the exception of the start codon, ranked [19]. Horizontal dotted lines represent putative cutoffs for defining viable mutations.

## Notes

https://github.com/allmanbrent/NLP_viral_escape

## References

[1] C. Chakraborty, A. R. Sharma, M. Bhattacharya, and S. S. Lee. A detailed overview of immune escape, antibody escape, partial vaccine escape of SARS-CoV-2 and their emerging variants with escape mutations. Front. Immunol., 13:801522, 2022. doi: 10.3389/fimmu.2022.801522.

[2] D. Focosi, M. Tuccori, A. Baj, and F. Maggi. SARS-CoV-2 variants: A synopsis of in vitro efficacy data of convalescent plasma, currently marketed vaccines, and monoclonal antibodies. Viruses, 13:1211, 2021. doi: 10.3390/v13071211.

[3] J. Zhou, K. Sukhova, T. P. Peacock, P. F. McKay, J. C. Brown, R. Frise, L. Baillon, M. Moshe, R. Kugathasan, R. J. Shattock, and W. S. Barclay. Omicron breakthrough infections in vaccinated or previously infected hamsters. Proc. Natl. Acad. Sci., 120: e2308655120, 2023. doi: 10.1073/PNAS.2308655120.

[4] M. B. Doud and J. D. Bloom. Accurate measurement of the effects of all amino-acid mutations on influenza hemagglutinin. Viruses, 8:155, 2016. doi: 10.3390/v8060155.

[5] A. Moulana, T. Dupic, A. M. Phillips, J. Chang, S. Nieves, A. A. Roffler, A. J. Greaney, T. N. Starr, J. D. Bloom, and M. M. Desai. Compensatory epistasis maintains ACE2 affinity in SARS-CoV-2 Omicron BA.1. Nat. Commun., 13:7011, 2022. doi: 10.1038/s41467-022-34506-z.

[6] A. Moulana, T. Dupic, A. M. Phillips, J. Chang, A. A. Roffler, A. J. Greaney, T. N. Starr, J. D. Bloom, and M. M. Desai. The landscape of antibody binding affinity in SARS-CoV-2 Omicron BA.1 evolution. eLife, 12:e83442, 2023. doi: 10.7554/eLife.83442.

[7] T. N. Starr, A. J. Greaney, A. Addetia, W. W. Hannon, M. C. Choudhary, A. S. Dingens, J. Z. Li, and J. D. Bloom. Prospective mapping of viral mutations that escape antibodies used to treat COVID-19. Science, 371:850–854, 2021. doi: 10.1126/science.abf9302.

[8] F. Obermeyer, M. Jankowiak, N. Barkas, S. F. Schaffner, J. D. Pyle, L. Yurkovetskiy, M. Bosso, D. J. Park, M. Babadi, B. L. MacInnis, J. Luban, P. C. Sabeti, and J. E. Lemieux. Analysis of 6.4 million SARS-CoV-2 genomes identifies mutations associated with fitness. Science, 376:1327–1332, 2022. doi: 10.1126/science.abm1208.

[9] D. Wang, M. Huot, V. Mohanty, and E. I. Shakhnovich. Biophysical principles predict fitness of SARS-CoV-2 variants. Proc. Natl. Acad. Sci., 121:e2314518121, 2024. doi: 10.1073/PNAS.2314518121.

[10] B. Hie, E. D. Zhong, B. Berger, and B. Bryson. Learning the language of viral evolution and escape. Science, 371:284–288, 2021. doi: 10.1126/science.abd7331.

[11] K. D. Lamb, J. Hughes, S. Lytras, O. Koci, F. Young, J. Grove, K. Yuan, and D. L. Robertson. From a single sequence to evolutionary trajectories: protein language models capture the evolutionary potential of SARS-CoV-2 protein sequences. bioRxiv, 2024. doi: 10.1101/2024.07.05.602129.

[12] N. Brandes, D. Ofer, Y. Peleg, N. Rappoport, and M. Linial. ProteinBERT: A universal deep-learning model of protein sequence and function. Bioinformatics, 38:2102–2110, 2022. doi: 10.1093/bioinformatics/btac020.

[13] Z. Lin, H. Akin, R. Rao, B. Hie, Z. Zhu, W. Lu, N. Smetanin, R. Verkuil, O. Kabeli, Y. Shmueli, A. dos Santos Costa, M. Fazel-Zarandi, T. Sercu, S. Candido, and A. Rives. Evolutionary-scale prediction of atomic-level protein structure with a language model. Science, 379:1123–1130, 2023. doi: 10.1126/science.ade2574.

[14] A. Madani, B. Krause, E. R. Greene, S. Subramanian, B. P. Mohr, J. M. Holton, J. L. Olmos, C. Xiong, Z. Z. Sun, R. Socher, J. S. Fraser, and N. Naik. Large language models generate functional protein sequences across diverse families. Nat. Biotechnol., 41:1099–1106, 2023. doi: 10.1038/s41587-022-01618-2.

[15] J. H. Lau, A. Clark, and S. Lappin. Grammaticality, acceptability, and probability: A probabilistic view of linguistic knowledge. Cogn. Sci., 41:1202–1241, 2017. doi: 10.1111/cogs.12414.

[16] P. D. Turney and P. Pantel. From frequency to meaning: Vector space models of semantics. *J*. Artif. Intell., 37:141–188, 2010. doi: 10.1613/jair.2934.

[17] M. B. Doud, J. M. Lee, and J. D. Bloom. How single mutations affect viral escape from broad and narrow antibodies to H1 influenza hemagglutinin. Nat. Commun., 9:1386, 2018. doi: 10.1038/s41467-018-03665-3.

[18] A. S. Dingens, D. Arenz, H. Weight, J. Overbaugh, and J. D. Bloom. An antigenic atlas of HIV-1 escape from broadly neutralizing antibodies distinguishes functional and structural epitopes. Immunity, 50:520–532.e3, 2019. doi: 10.1016/j.immuni.2018.12.017.

[19] H. K. Haddox, A. S. Dingens, S. K. Hilton, J. Overbaugh, and J. D. Bloom. Mapping mutational effects along the evolutionary landscape of HIV envelope. eLife, 7:e34420, 2018. doi: 10.7554/eLife.34420.

[20] A. Baum, B. O. Fulton, E. Wloga, R. Copin, K. E. Pascal, V. Russo, S. Giordano, K. Lanza, N. Negron, M. Ni, Y. Wei, G. S. Atwal, A. J. Murphy, N. Stahl, G. D. Yancopoulos, and C. A. Kyratsous. Antibody cocktail to SARS-CoV-2 spike protein prevents rapid mutational escape seen with individual antibodies. Science, 369, 2020. doi: 10.1126/science.abd0831.

[21] S. d’Oelsnitz, D. J. Diaz, W. Kim, D. J. Acosta, T. L. Dangerfield, M. W. Schechter, M. B. Minus, J. R. Howard, H. Do, J. M. Loy, H. S. Alper, Y. J. Zhang, and A. D. Ellington. Biosensor and machine learning-aided engineering of an amaryllidaceae enzyme. Nat. Commun., 15:2084, 2024. doi: 10.1038/s41467-024-46356-y.

[22] C. Gong, A. Klivans, J. M. Loy, T. Chen, Q. Liu, and D. J. Diaz. Evolution-inspired loss functions for protein representation learning. Proc. Int. Conf. Mach. Learn., 2024.

[23] A. V. Kulikova, D. J. Diaz, T. Chen, T. J. Cole, A. D. Ellington, and C. O. Wilke. Two sequence-and two structure-based ML models have learned different aspects of protein biochemistry. Sci. Rep., 13:13280, 2023. doi: 10.1038/s41598-023-40247-w.

[24] W. Torng and R. B. Altman. 3D deep convolutional neural networks for amino acid environment similarity analysis. BMC Bioinform., 18:302, 2017. doi: 10.1186/s12859-017-1702-0.

[25] D. J. Diaz, A. V. Kulikova, A. D. Ellington, and C. O. Wilke. Using machine learning to predict the effects and consequences of mutations in proteins. Curr. Opin. Struct. Biol., 78:102518, 2023. doi: 10.1016/j.sbi.2022.102518.

[26] A. Elnaggar, M. Heinzinger, C. Dallago, G. Rehawi, Y. Wang, L. Jones, T. Gibbs, T. Feher, C. Angerer, M. Steinegger, D. Bhowmik, and B. Rost. ProtTrans: Toward understanding the language of life through self-supervised learning. IEEE Trans. Pattern Anal. Mach. Intell., 44:7112–7127, 2022. doi: 10.1109/TPAMI.2021.3095381.

[27] Y. Geffen, Y. Ofran, and R. Unger. DistilProtBert: a distilled protein language model used to distinguish between real proteins and their randomly shuffled counterparts. Bioinformatics, 38:ii95–ii98, 2022. doi: 10.1093/bioinformatics/btac474.

[28] E. Capriotti, P. Fariselli, I. Rossi, and R. Casadio. A three-state prediction of single point mutations on protein stability changes. BMC Bioinform., 9:S6, 2008. doi: 10.1186/1471-2105-9-S2-S6.

[29] B. E. Suzek, Y. Wang, H. Huang, P. B. McGarvey, C.H. Wu, and the UniProt Consortium. UniRef clusters: a comprehensive and scalable alternative for improving sequence similarity searches. Bioinformatics, 31:926–932, 2015. doi: 10.1093/bioinformatics/btu739.

[30] D. J. Diaz, C. Gong, J. Ouyang-Zhang, J. M. Loy, J. Wells, D. Yang, A. D. Ellington, A. G. Dimakis, and A. R. Klivans. Stability oracle: A structure-based graph-transformer for identifying stabilizing mutations. Nat. Commun., 15:6170, 2024. doi: 10.1038/s41467-024-49780-2.

[31] K. Tsuboyama, J. Dauparas, J. Chen, E. Laine, Y. Mohseni Behbahani, J. J. Weinstein, N. M. Mangan, S. Ovchinnikov, and G. J. Rocklin. Mega-scale experimental analysis of protein folding stability in biology and design. Nature, 620(7973):434–444, August 2023. ISSN 1476-4687. doi: 10.1038/s41586-023-06328-6.

[32] J. Devlin, M. Chang, K. Lee, and K. Toutanova. BERT: Pre-training of deep bidirectional transformers for language understanding. arXiv, 2019. doi: 10.48550/arXiv.1810.04805.

[33] R. Shroff, A. W. Cole, D. J. Diaz, B. R. Morrow, I. Donnell, A. Annapareddy, J. Gollihar, A. D. Ellington, and R. Thyer. Discovery of novel gain-of-function mutations guided by structure-based deep learning. ACS Synth. Biol., 9:2927–2935, 2020. doi: 10.1021/acssynbio.0c00345.

[34] I. Paik, P. H. T. Ngo, R. Shroff, D. J. Diaz, A. C. Maranhao, D. J. F. Walker, S. Bhadra, and A. D. Ellington. Improved bst dna polymerase variants derived via a machine learning approach. Biochemistry, 2021. doi: 10.1021/acs.biochem.1c00451.

[35] H. Lu, D. J. Diaz, N. J. Czarnecki, C. Zhu, W. Kim, R. Shroff, D. J. Acosta, B. R. Alexander, H. O. Cole, Y. Zhang, et al. Machine learning-aided engineering of hydrolases for PET depolymerization. Nature, 604:662–667, 2022. doi: 10.1038/s41586-022-04599-z.

[36] Y. Liu, S. G. Bender, D. Sorigue, D. J. Diaz, A. D. Ellington, G. Mann, S. Allmendinger, and T. K. Hyster. Asymmetric synthesis of *α*-chloroamides via photoenzymatic hydroalkylation of olefins. J. Am. Chem. Soc., 146:7191–7197, 2024. doi: 10.1021/jacs.4c00927.

[37] R. Hunter Wilson, Anoop R. Damodaran, and Ambika Bhagi-Damodaran. Machine learning guided rational design of a non-heme iron-based lysine dioxygenase improves its total turnover number. bioRxiv, 2024. doi: 10.1101/2024.06.04.597480.

[38] J. D. Bloom, J. J. Silberg, C. O. Wilke, D. A. Drummond, C. Adami, and F. H. Arnold. Thermodynamic prediction of protein neutrality. Proc. Natl. Acad. Sci., 102:606–611, 2005. doi: 10.1073/PNAS.0406744102.

[39] L. I. Gong, M. A. Suchard, and J. D. Bloom. Stability-mediated epistasis constrains the evolution of an influenza protein. eLife, 2:e00631, 2013. doi: 10.7554/eLife.00631.

[40] D. A. Liberles, S. A. Teichmann, I. Bahar, U. Bastolla, J. Bloom, E. Bornberg-Bauer, L. J. Colwell, A. P. J. de Koning, N. V. Dokholyan, J. Echave, A. Elofsson, D. L. Gerloff, R. A. Goldstein, J. A. Grahnen, M. T. Holder, C. Lakner, N. Lartillot, S. C. Lovell, G. Naylor, T. Perica, D. D. Pollock, T. Pupko, L. Regan, A. Roger, N. Rubinstein, E. Shakhnovich, K. Sjőlander, S. Sunyaev, A. I. Teufel, J. L. Thorne, J. W. Thornton, D. M. Weinreich, and S. Whelan. The interface of protein structure, protein biophysics, and molecular evolution. Protein Sci., 21:769–785, 2012. doi: 10.1002/pro.2071.

[41] A. Rotem, A. W. R. Serohijos, C. B. Chang, J. T. Wolfe, A. E. Fischer, T. S. Mehoke, H. Zhang, Y. Tao, W. Lloyd Ung, J.-M. Choi, J. V. Rodrigues, A. O. Kolawole, S. A. Koehler, S. Wu, P. M. Thielen, N. Cui, P. A. Demirev, N. S. Giacobbi, T. R. Julian, K. Schwab, J. S. Lin, T. J. Smith, J. M. Pipas, C. E. Wobus, A. B. Feldman, D. A. Weitz, and E. I. Shakhnovich. Evolution on the biophysical fitness landscape of an RNA virus. Mol. Biol. and Evol., 35:2390–2400, 2018. doi: 10.1093/molbev/msy131.

[42] C. Scott Wylie and Eugene I. Shakhnovich. A biophysical protein folding model accounts for most mutational fitness effects in viruses. Proc. Natl. Acad. Sci., 108:9916–9921, 2011. doi: 10.1073/PNAS.1017572108.

[43] J. Schymkowitz, J. Borg, F. Stricher, R. Nys, F. Rousseau, and L. Serrano. The FoldX web server: an online force field. Nucleic Acids Res., 33:W382–W388, 2005. doi: 10.1093/nar/gki387.

[44] Y. Dehouck, J. M. Kwasigroch, D. Gilis, and M. Rooman. PoPMuSiC 2.1: A web server for the estimation of protein stability changes upon mutation and sequence optimality. BMC Bioinform., 12:151, 2011. doi: 10.1186/1471-2105-12-151.

[45] V. Sora, A. O. Laspiur, K. Degn, M. Arnaudi, M. Utichi, L. Beltrame, D. De Menezes, M. Orlandi, U. K. Stoltze, O. Rigina, P. W. Sackett, K. Wadt, K. Schmiegelow, M. Tiberti, and E. Papaleo. RosettaDDGPrediction for high-throughput mutational scans: From stability to binding. Protein Sci., 32:e4527, 2023. doi: 10.1002/pro.4527.

[46] C. O. Wilke. The biophysical landscape of viral evolution. Proc. Natl. Acad. Sci., 121: e2409667121, 2024. doi: 10.1073/PNAS.240966712.

[47] M. Littmann, M. Heinzinger, C. Dallago, T. Olenyi, and B. Rost. Embeddings from deep learning transfer GO annotations beyond homology. Sci. Rep., 2021. doi: 10.1038/s41598-020-80786-0.

[48] M. Heinzinger, A. Elnaggar, Y. Wang, C. Dallago, D. Nechaev, F. Matthes, and B. Rost. Modeling aspects of the language of life through transfer-learning protein sequences. BMC Bioinform., 20:723, 2019. doi: 10.1186/s12859-019-3220-8.

[49] J. R. Randall, L. C. Vieira, C. O. Wilke, and B. W. Davies. Deep mutational scanning and machine learning for the analysis of antimicrobial-peptide features driving membrane selectivity. *Nat*. Biomed. Eng, 2024. doi: 10.21203/rs.3.rs-3280212/v1.

[50] A. Villegas-Morcillo, S. Makrodimitris, R. C. H. J. van Ham, A. M. Gomez, V. Sanchez, and M. J. T. Reinders. Unsupervised protein embeddings outperform hand-crafted sequence and structure features at predicting molecular function. Bioinformatics, 2021. doi: 10.1093/bioinformatics/btaa701.

[51] J. Ouyang-Zhang, D. Diaz, A. Klivans, and P. Kraehenbuehl. Predicting a protein’s stability under a million mutations. Adv. Neural. Inf. Process. Syst., 36:76229–76247, 2023.

[52] M. Kulmanov, F. J. Guzmán-Vega, P. Duek Roggli, L. Lane, S. T. Arold, and R. Hoehndorf. Protein function prediction as approximate semantic entailment. Nat. Mach. Intell., 6:220–228, 2024. doi: 10.1038/s42256-024-00795-w.

[53] C. Gong, A. Klivans, J. Wells, J. Loy, Q. Liu, A. Dimakis, and D. Diaz. Binding oracle: Fine-tuning from stability to binding free energy. NeurIPS 2023 Generative AI and Biology (GenBio) Workshop, 2023. URL https://openreview.net/forum?id=ChU7MCLk1J.

[54] R. Evans, M. O’Neill, A. Pritzel, N. Antropova, A. Senior, T. Green, A. Žídek, R. Bates, S. Blackwell, J. Yim, O. Ronneberger, S. Bodenstein, M. Zielinski, A. Bridgland, A. Potapenko, A. Cowie, K. Tunyasuvunakool, R. Jain, E. Clancy, P. Kohli, J. Jumper, and D. Hassabis. Protein complex prediction with AlphaFold-Multimer. bioRxiv, 2022. doi: 10.1101/2021.10.04.463034v2.

[55] J. Jumper, R. Evans, A. Pritzel, T. Green, M. Figurnov, O. Ronneberger, K. Tunyasuvunakool, R. Bates, A. Žídek, A. Potapenko, A. Bridgland, C. Meyer, S. A. A. Kohl, A. J. Ballard, A. Cowie, B. Romera-Paredes, S. Nikolov, R. Jain, J. Adler, T. Back, S. Petersen, D. Reiman, E. Clancy, M. Zielinski, M. Steinegger, M. Pacholska, T. Berghammer, S. Bodenstein, D. Silver, O. Vinyals, A. W. Senior, K. Kavukcuoglu, P. Kohli, and D. Hassabis. Highly accurate protein structure prediction with AlphaFold. Nature, 596: 583–589, 2021. doi: 10.1038/s41586-021-03819-2.

[56] J. D. Bloom. Software for the analysis and visualization of deep mutational scanning data. BMC Bioinform., 16:168, 2015. doi: 10.1186/s12859-015-0590-4.

